# HMMerge: an Ensemble Method for Improving Multiple Sequence Alignment

**DOI:** 10.1101/2022.05.29.493880

**Authors:** Minhyuk Park, Tandy Warnow

## Abstract

Despite advances in method development for multiple sequence alignment over the last several decades, the alignment of datasets exhibiting substantial sequence length heterogeneity, especially when the input sequences include very short sequences (either as a result of sequencing technologies or of large deletions during evolution) remains an inadequately solved problem. We present HMMerge, a method to compute an alignment of datasets exhibiting high sequence length heterogeneity, or to add short sequences into a given “backbone” alignment. HMMerge builds on the technique from its predecessor alignment methods, UPP and WITCH, which build an ensemble of HMMs for the backbone alignment and add the remaining sequences into the backbone alignment using the ensemble. HMMerge differs from UPP and WITCH by building a new HMM for each query sequence: it uses a novel ensemble approach to combine the HMMs, each weighted by the probability of generating the query sequence, into a single HMM. Then it applies the Viterbi algorithm to add the query sequence into the backbone alignment. We show that using this “merged” HMM provides better accuracy than the current approach in UPP and matches or improves on WITCH for adding short sequences into backbone alignments. HMMerge is freely available at https://github.com/MinhyukPark/HMMerge.

## 1 Introduction

Multiple sequence alignment (MSA) is a necessary first step in many common bioinformatics pipelines, such as phylogenetic tree inference and metagenomic taxon identification, and so obtaining high quality MSAs has high relevance in downstream analyses [8, 11]. One of the challenges in multiple sequence alignment is when the input sequence dataset has highly variable sequence lengths, a property that is found in many biological datasets [12]. Sequence length heterogeneity can be caused by evolutionary processes (i.e., large indels), but is also produced when datasets include reads generated by Illumina and other short read sequencing technologies [15].

UPP [12] and its successor WITCH [14] are methods that are designed to align datasets when the input has sequence length heterogeneity. Both methods use a two-stage approach, where they first identify and extract a subset of the input sequences (based on their sequence length) that are considered to be “fulllength”, and then align these sequences, forming the “backbone alignment”. Both then builds an ensemble of profile Hidden Markov Models (eHMM) to represent the backbone alignment, where each profile HMM is built on a subset of the input sequences in the backbone alignment. UPP uses a straightforward method to add each of the remaining sequences (which are called “query sequences) into the backbone alignment: each query sequence (a) selects a best-fitting HMM from the eHMM based on the bit scores, (b) the Viterbi algorithm is used to find an optimal path for the query sequence to the selected HMM, and (c) the match states in the optimal path define an alignment of the query sequence to the backbone alignment, which is used to add the query sequence into the backbone alignment. WITCH elaborates on this approach by weighting each profile HMM with the probability it has of generating the query sequence, and then combines the optimal alignments (one such alignment for each profile HMM in eHMM) using a weighted consensus algorithm that in turn is based on the Graph Clustering Merger (GCM) method from the alignment method MAGUS [17]. Both methods produce more accurate alignments when datasets have fragmentary sequences [14, 12], but WITCH produces more accurate alignments than UPP [14]. We present a sample of results comparing UPP, WITCH, and other alignment methods (Table 1) on 1000-sequence simulated datasets with varying rates of evolution, each having large numbers of short sequences, demonstrating that UPP and WITCH both produce more accurate alignments than the other methods.

**Table 1:** Sequence alignment error on simulated datasets with introduced fragmentation. We show the average of SPFN error (proportion of missing homologies) and SPFP error (proportion of false homologies) for alignments produced by UPP, WITCH, and leading MSA methods on simulated datasets with 1000 sequences, high rates of evolution, and varying gap lengths (L for long, M for medium, and S for short). All datasets have high fragmentation (50% of the sequences are fragmentary). See Supplementary Materials for additional information.

|  | 1000L1-HF | 1000M1-HF | 1000S1-HF |
| --- | --- | --- | --- |
| WITCH | **0.086** | **0.149** | **0.130** |
| UPP | 0.093 | 0.155 | 0.136 |
| MAGUS | 0.199 | 0.289 | 0.271 |
| PASTA | 0.480 | 0.647 | 0.590 |
| MAFFT | 0.480 | 0.583 | 0.581 |
| Clustal Omega | 0.935 | 0.951 | 0.929 |
| T-COFFEE | 0.953 | 0.961 | 0.929 |
| MUSCLE | 0.635 | 0.816 | 0.781 |

**Table 2:** Biological DNA/RNA Dataset Overview. Here, we show the basic empirical statistics about the datasets used in this study. p-dist. refers to the normalized Hamming distance, which is computed by dividing the number of positions where two sequences are both non-gapped and have different characters by the number of positions where two sequences are non-gapped.

| Name | # Sequences | avg. p-dist. | avg len. |
| --- | --- | --- | --- |
| 23S.A | 214 | 0.293 | 1851 |
| 23S.C | 374 | 0.143 | 2087 |
| 5S.3 | 5507 | 0.417 | 106 |
| 5S.E | 2774 | 0.305 | 96 |
| 5S.T | 5751 | 0.425 | 106 |

In this study, we present HMMerge, a new approach for adding sequences into backbone alignments. We follow the same initial steps as WITCH and UPP (i.e., we use the same technique to build the backbone alignment and the ensemble of HMMs representing the backbone alignment). Given a query sequence, we use the same weighting scheme as in WITCH to weight each HMM based on the probability that the given HMM generates the given query sequence. For that given query sequence, we then build a new HMM that is the weighted combination of the HMMs in the eHMM. Applying the Viterbi algorithm to the query sequence and the HMM defines an optimal alignment of the query sequence to the backbone alignment. As this study shows, using both simulated and biological data, HMMerge matches or improves on the accuracy of WITCH, with specific advantages when the dataset is difficult to align due to evolving under a high rate of evolution.

## 2 Background

Here we present UPP and WITCH, which are the two methods most closely related to HMMerge. UPP (Ultra-large alignment using Phylogenetic Profiles) and WITCH (WeIghTed Consensus Hmm alignment) are methods that are designed to align sequence datasets that contain substantial sequence length heterogeneity. When aligning datasets with many short sequences, WITCH and UPP both produce more accurate alignments than standard alignment methods. A sample of these results on three model conditions is shown in Table 1, each with 1000 sequences and high rates of evolution, a large fraction of fragmentary sequences, and three gap lengths (long, medium, and short). Note that WITCH is more accurate than UPP, but UPP is also more accurate than all the remaining methods. Additional results on other model conditions are shown in the Supplementary Materials, and show that these trends hold under lower rates of evolution, but with smaller gaps between methods.

Both UPP and WITCH operate in three stages: the backbone stage, the construction of the eHMM for the backbone, and then the search-and-align stage. UPP and WITCH are identical for the first two stages but differ in the search-and-align stage.

### Stage 1: Computing the Backbone Alignment

In the backbone stage, the input sequences are split into two sets, with one set containing some of the “full-length” sequences and the remaining sequences denoted “query” sequences. An alignment is built on the full-length sequences, using external methods (e.g., PASTA [9] or MAGUS [17].

### Stage 2: Computing the eHMM

Once the backbone alignment is computed, a tree is computed on the backbone alignment (e.g., using a maximum likelihood method FastTree 2 [13] or RAxML [19]). Then, the backbone tree is decomposed into subsets by edge deletions (removing “centroid edges” that roughly split the leaf-set in half) until each subset is not too large (the default in UPP is to decompose until each subset has at most 10 leaves). The full set of backbone sequences and any subset produced by the decomposition pipeline becomes one of the subsets. A profile HMM is constructed on each subset using *hmmbuild* from the HMMER suite [3]. The collection of profile HMMs produced in this stage is referred to as an “ensemble of HMMs” for the backbone alignment, or “eHMM”.

### Stage 3: Search-and-align stage

Since WITCH improves on UPP and they differ only in this third stage, we describe only how WITCH performs this stage. To add a given query sequence into the backbone alignment using WITCH, each query sequence scores each HMM in the eHMM according to an “adjusted” bit score, which is designed to be an estimate of the probability that the HMM generated the given query sequence (see [14] for the derivation of the adjusted bit score). For each HMM in the eHMM, WITCH uses hmmalign (from HMMER [3]) to compute an alignment of the query sequence to the induced subset alignment, and hence into the backbone alignment. Thus, each query sequence is added into the backbone alignment in multiple ways. These different extended alignments are weighted by the adjusted bit scores, and then combined into a single extended alignment (that includes the query sequence and the backbone alignment) using a weighted version of the Graph Clustering Merger from MAGUS.

Thus, the search-and-align stage adds each query sequence into the backbone alignment. The final merged alignment of all the query sequences is done using transitivity.

#### UPP-add and WITCH-add

The second and third stages for UPP and WITCH define subroutines that take as input the backbone alignment and add the remaining sequences into the backbone alignment. Thus, we can refer to these methods, used just for these two steps, as UPP-add and WITCH-add.

## 3 Algorithm: HMMerge

HMMerge has the same three-stage procedure as UPP and WITCH and differs from them also only in the third stage. Thus, we will refer to HMMerge for the pipeline, and HMMerge-add for the use of HMMerge when the backbone alignment is already given.

### 3.1 High-level description of HMMerge

We now describe at a high level how HMMerge operates, given the input backbone alignment *A* and query sequence set *Q*. We follow the same steps as UPP and WITCH for Stage 1 (Backbone Alignment) and Stage 2 (Building the eHMM for the backbone alignment). Hence, we only need to define how we run Stage 3 (Search-and-align), which is to say how we run HMMerge-add.

We describe HMMerge-add in its most general context, where the input is a multiple sequence alignment (which we will refer to as the backbone alignment) and additional sequences (called query sequences) and the output is an alignment on the full set of sequences (backbone sequences and query sequences) that induces the backbone alignment. Thus, the problem that the HMMerge-add addresses is as follows:

- Input: Backbone alignment *A* and query sequence set *Q*
- Output: Extended alignment *A*\* that induces *A* and includes all the sequences in *Q*

We now provide details for how HMMerge operates, beginning with terminology we will use.

### 3.2 Terminology

#### substring

Given a sequence, or string, *S*, *S*[*i*] denotes the ith character in the sequence while *S*[*i*..*j*] denotes the substring starting at character *S*[*i*] and ending at character *S*[*j*]. The first letter is *S*[1] throughout this document.

#### Indexing

Given an indexable object such as a matrix, index · refers to the entire range of the current axis. For example, *A*[·] refers to all entries of *A* while *B*[1, ·] refers to all column entries of B whose row index is 1.

#### Precondition

The adjacency matrix *M*, transition probability matrix *T*, and emission probability matrix *P* encode the states of an HMM. These states must be ordered such that for any two indices where *i* < *j*, state *i* appears before state *j* in the matrix ordering, and that state *i* appears before state *j* in a topologically sorted graph that is represented by *M*.

- Σ is the set of letters that can be emitted. The set Σ is a finite set.
- *n* is the number of states in our HMM
- *M*, the adjacency matrix of dimensions n by *n*, which represents the HMM. The entry *M*[*i*, *j*] is 1 if there is an edge from state *i* to state *j* and *i* ≠ *j*. If there is an edge from state *i* to state *i*, then the entry *M*[*i*, *i*] is 2. If there are no edges from state *i* to state *j*, then *M*[*i*, *j*] is 0. Although not shown in formulas, this matrix is used internally to track the edge information as the final HMM gets updated.

— *M*[*i*,*j*] = 0 if there is no edge from state *i* to state *j*
— *M* [*i*,*j*] = 1 if there is an edge from state *i* to state *j*
— *M*[*i*, *i*] = 2 if there is an edge from state *i* to state *i*
- *T*, the transition probability matrix with the same dimensions as *M* where *T*[*i*,*j*] indicates the transition probability of going from state *i* to state *j*. If there are no edges from state i to state *j*, then the entry *T* [*i*,*j*] is 0. Since the matrix *T* denotes the transition probabilities, the row sum of any entry is 1 except for the final accepting state, which does not have any outgoing edges and hence has a row sum of 0.
- *P*, the emission probability matrix of dimensions *n* by |Σ| where *P*[*i*, *s*] is the probability of state *i* emitting letter *s*. If state i does not emit any characters, then *P*[*i*, ·] = 0. If state *i* is an emission state, *P*[*i*, *ϵ*] = 0 where ε is the empty string.

### 3.3 Merged HMM Construction

Here, we describe the procedure for obtaining a hidden Markov model from a set of input profile hidden Markov models and a query sequence.

#### 1. Topology Construction

For every column in the backbone alignment, we need to identify which HMMs provide relevant information. Once the relevant HMMs are identified, all of their edges are inserted into the output HMM. To define relevant HMMs in this context, since subset alignments will have fully gapped columns removed, it is possible for certain HMMs to entirely omit columns that exist in the backbone alignment. Therefore, not all HMMs will provide information about every column in the backbone alignment. In those cases where backbone columns are omitted in the subset alignment, the backbone column immediately before and after the omitted columns are considered to be connected. This is represented with transition edges that skip over omitted columns in the final output HMM.

#### 2. Numeric Parameters

Although the topology remains fixed for each query sequence, the numeric parameters vary for each query sequence. The weights of each HMM, and hence the weights of the edges each HMM contributes, are calculated according to the adjusted bit-score of the query sequence to each of the HMMs. This is done through the derivation first presented in [14]. The transition probabilities, insertion probabilities, and emission probabilities from each HMM are averaged with their weights derived from the adjusted bit-scores.

The result is a hidden Markov model that contains all the edges from the relevant input profile hidden Markov models with the transition, insertion, and emission probabilities weighted according to the adjusted bit-scores. Here, *n* is always equal to the number of columns in the backbone alignment.

### 3.4 Matrix Representation

In order to represent an HMM with matrices, we need a mapping from the states to a matrix index. Suppose that |*B*| is the number of columns in the backbone alignment. Since every column has a match state, insertion state, and a deletion state, the HMM has 3|*B*| states that are associated with a backbone alignment column. In addition, there are start and end states along with an insertion state connected to the start state.

In total, there are 3|*B*| + 3 number of states in our HMM. See Figure 1 for an example HMM. We can map the states in such a way where the ordering is [start, *I*_0_, *M*_1_, *I*_1_, *D*_1_, *M*_2_, *I*_2_, *D*_2_,…, *M*_|*B*|_, *I*_|*B*|_, *D*_|*B*|_, end]. For the match state corresponding to the ith column (*M_i_*), the index in the ordering is 2 + 3(*i* – 1). For the insertion state corresponding to the ith column (*I_i_*), the index in the ordering is 2 +3(*i* – 1) + 1. For the deletion state corresponding to the ith column (*D_i_*), the index in the ordering is 2 + 3(*i* – 1) + 2. This ordering ensures that a state that comes before another state in the HMM will also come before the other state in the numeric ordering, thus topologically sorting the HMM.

**Figure 1:**
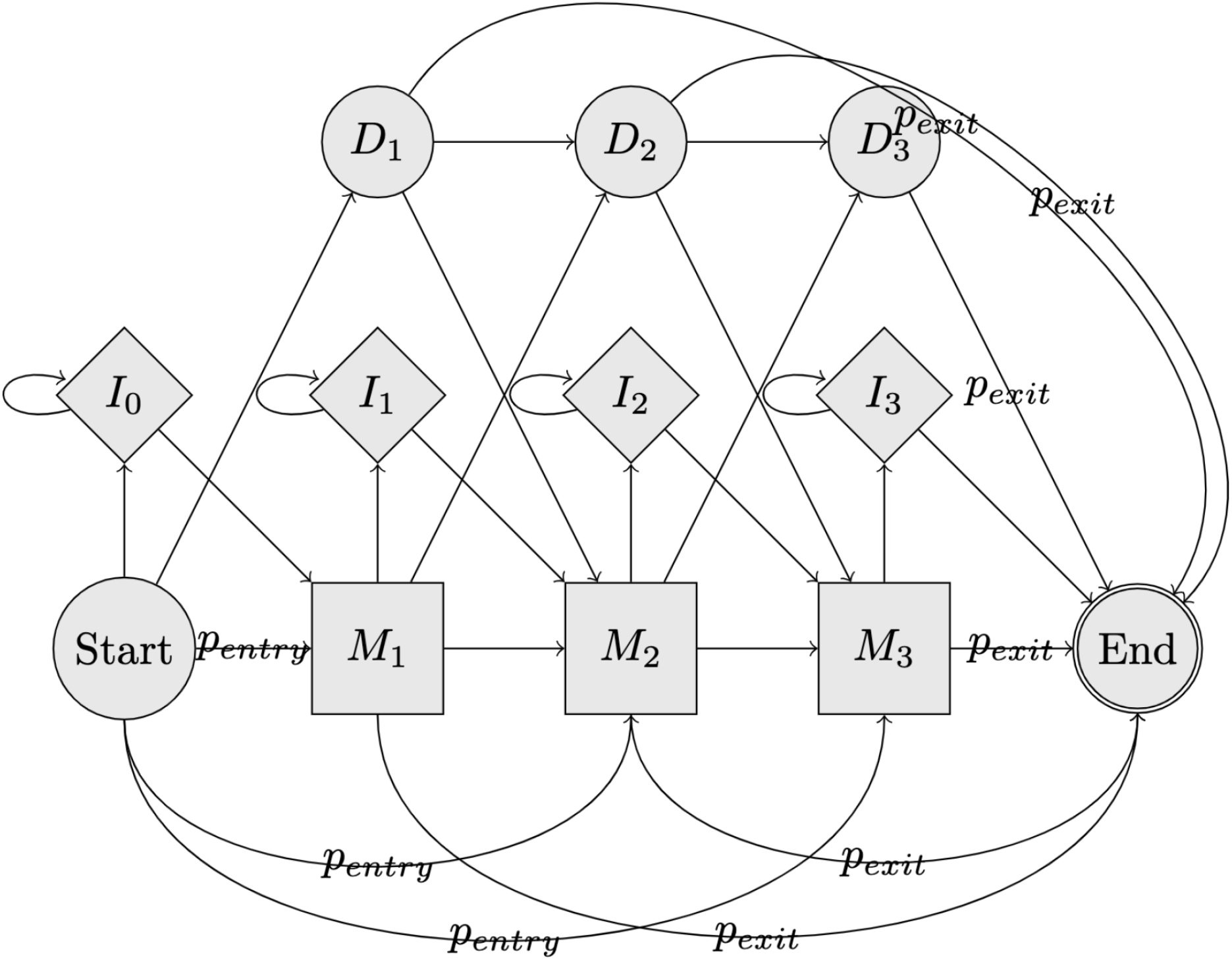
Equal Entry Exit HMM Example. Here, we show a visualization of an example HMM with equal entry and exit probability edges. *P_entry_* and *P_exit_* represent the probabilities of a path taking that edge. *M_i_* stands for the ith match state, *I*_i_ stands for the ith insertion state, and *D*_i_ stands for the ith deletion state. The start and end states do not emit any characters.

### 3.5 Alignment of Sequence to Merged HMM

Here, we describe global alignment and glocal alignment. We will also explain how global and glocal alignment are implemented in this paper.

#### 3.5.1 Global alignment (Viterbi Algorithm)

Given two sequences, a global alignment is a type of alignment that aligns the ends of one sequence to the ends of another sequence. In other words, global alignment aligns the sequences “end-to-end”. This is implemented via the Viterbi algorithm. We followed the standard Viterbi algorithm with additional cares to ensure that states that do not emit characters are handled properly.

##### Inputs

We define a table VT for ViTerbi. Suppose the input sequence *S* has length *L*. The parameters to our function are *i* and *k* where VT[i,k] represents the maximum probability of the path that emits the letters *S*[0..*i*] and ending at state *k*. Given a matrix where the rows correspond to the states *k* and the columns correspond to the indices *i*, since every VT entry only depends on VT entries where either the row index is less than *i*, or the column index is less than *k*, the evaluation order can simply be a column-major ordering. We can start filling in the entries of our table by starting at the first column and going down the rows.

###### Base cases

- *VT*[0, 0] = 1 Here, we are assuming that 0 is the index of the start state. This is the top left corner of the matrix if we imagine the VT table to have row indices going from top to bottom and the column indices going from left to right. This base case and the first case in the subproblem together fill out the first row of the VT table.
- *VT*[0, *j*] = 0 if *j* is an emission state. This base case and the second case in the subproblem together fill out the first column of the VT table.

###### Subproblems

State *j* is an emission state if Σ_*k*∈Σ_*P*[*j*, *k*] > 0

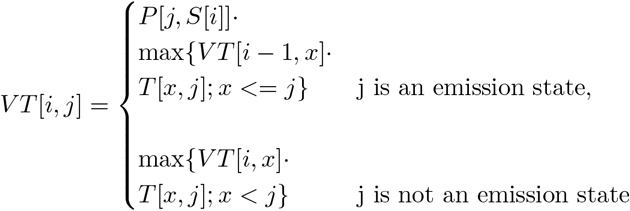

For all nodes that have an outgoing edge to our current state *j*, we multiply the probability of emitting the current character *S* [*i*] at our current state *j* with the maximum probability of ending up at our current state.

##### Log-Likelihoods

In order to save computation time and achieve better numerical stability, we implemented a log-likelihood version of the Viterbi algorithm described above. Now, VT[i,k] represents the maximum log probability of the path that emits the letters *S*[0..*i*] and ending at state *k*. We quickly redefine the base cases and the subproblems here.

###### Log-Likelihoods Base cases

- *VT*[0, 0] = 0 Since log_2_ 1 = 0.
- *VT*[0, *j*] = – inf if *j* is an emission state since log_2_ 0 = – inf. This base case and the second case in the subproblem together fill out the first column of the VT table.

###### Log-Likelihoods Subproblems

State *j* is an emission state if Σ_k∈Σ_*P*[*j*, *k*] > 0

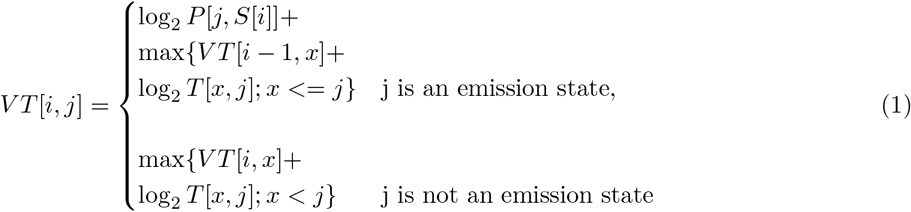

#### 3.5.2 Glocal alignment

A glocal alignment is a hybrid approach in which the entirety of one sequence is aligned to a segment from another sequence. An example where this may be useful is aligning a short sequence to a long sequence. In this case, we would like to align the entirety of the short sequence to a specific region in the longer sequence.

In order to allow for glocal alignment without incurring any penalties, i.e., aligning a query sequence to a specific region of the backbone alignment, we need to allow for our HMM algorithm to be able to start at any match state and end at any match state.

We now define two new probabilities, *p_entry_* and *p_ex_it,* which represent the transition probabilities from the start state to each match state and from each match and delete states to the end state, respectively. See Figure 1 for a visualization of how these edges are inserted. To account for the new edges, we re-normalize the edge weights such that all outgoing edge weights sum to 1 for each state. We describe this step in more detail in the Supplementary Materials.

### 3.6 Runtime Analysis

Supposed *S* is the number of sequences in the backbone alignment, |*B*| is the length of the backbone alignment, *d* is the decomposition size, and *q* is the number of query sequences. It takes *O*(*S*|*B*|) to parse the input sequences and create a mapping of the backbone alignment characters to input decomposed alignment characters. There are 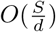 calls to *hmmbuild* to build the input HMMs and 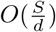 calls to *hmmsearch* to obtain the bit-scores of each query sequence for every HMM. For parsing outputs, it takes 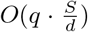 to parse the *hmmsearch* outputs and 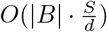 to read the *hmmbuild* outputs. Building the new HMMs takes 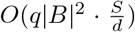 to calculate the new probabilities for the merged HMM, *O*(|*B*|^2^) to represent the HMM in probability matrices, and *O*(|*B*|_2_) to add new edges to enable glocal alignment. Finally, it takes *O*(*q*·|*B*|_3_) to run the Viterbi algorithm for each query sequence.

In total, the big O runtime is 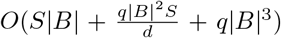excluding the 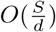 calls to *hmmbulid* and *hmmsearch*.

### 3.7 Experimental Study Design

It is possible to directly compare the effectiveness of HMMerge-add, WITCH-add, and UPP-add for adding query sequences given identical backbone alignments (and even identical eHMMs for the backbone align-ments). Our first experiment directly explored this on a variety of simulated datasets with introduced fragmentation. Our second experiment focused on nucleotide biological datasets.

#### Experiment 1: Comparison of HMMerge, WITCH, and UPP for adding query sequences on simulated datasets with introduced fragmentation

In Experiment 1, we explored the effectiveness of HMMerge, WITCH, and UPP for aligning query sequences to an existing backbone alignment on simulated datasets with introduced fragmentation. On all datasets tested, HMMerge, WITCH, and UPP used the same backbone alignments computed by MAGUS and backbone trees computed by FastTree [13]. In this experiment, we only looked at the alignment error when restricted to the query sequences.

#### Experiment 2: Comparison of HMMerge, WITCH, and UPP for adding query sequences on nucleotide biological datasets

In Experiment 2, we focused on biological datasets. On all datasets tested, HMMerge, WITCH, and UPP used the same backbone alignments computed by MAGUS and backbone trees computed by FastTree [13]. In this experiment, we only looked at the alignment error when restricted to the query sequences.

#### Datasets

We created high fragmentary versions of the ROSE simulated datasets, first studied in [7], using the same technique and parameters as used in [18], meaning that each sequence has been made fragmentary by taking half of the sequences and trimming them to be on average 25% of the median length with a standard deviation of 60 base pairs. These datasets consist of 500 full-length sequences and 500 fragmentary sequences. There are 15 different model conditions we explored, each with 20 replicates, where each model condition varies by the average length of gaps (“S” for Short, “M” for Medium, or “L” for Long) and by their varying evolutionary rates (numbered 1 through 5 the rate of evolution decreases as the number increases). Specifically, we used the high fragmentary versions of 1000S1-1000S5, 1000M1-1000M5, and 1000L1-1000L5.

We also used the high fragmentary, or “HF”, versions of the RNASim simulated datasets, taken from the study by [18], and made an ultra-high fragmentary, or “UHF”, version of the dataset by setting the target mean to be exactly half of the HF condition. We used 10 replicates each for both datasets. The target mean sequence length was 388 for the HF condition and 194 for the UHF condition. The standard deviation remained at 60, which is the same as the original study.

For biological nucleotide datasets, we used the CRW biological datasets from the Comparative RNA Website published by the Gutell lab [2]. Specifically, we chose 23S.A, 23S.C, 5S.3, 5S.E, and 5S.T for our study due to their sequence length distributions showing signs of high fragmentation (Figure 2). In order to split the datasets into backbone sequences and query sequences, we split the sequences manually by visual inspection of their sequence length histograms. As a result, 23S.A and 23S.C were both split at 1250, and 5S.3, 5S.E, and 5S.T were split at 100. That is to say that on the 23S datasets, any sequence with length greater than 1250 was chosen as the backbone while the remaining sequences were chosen to be query sequences.

**Figure 2:**
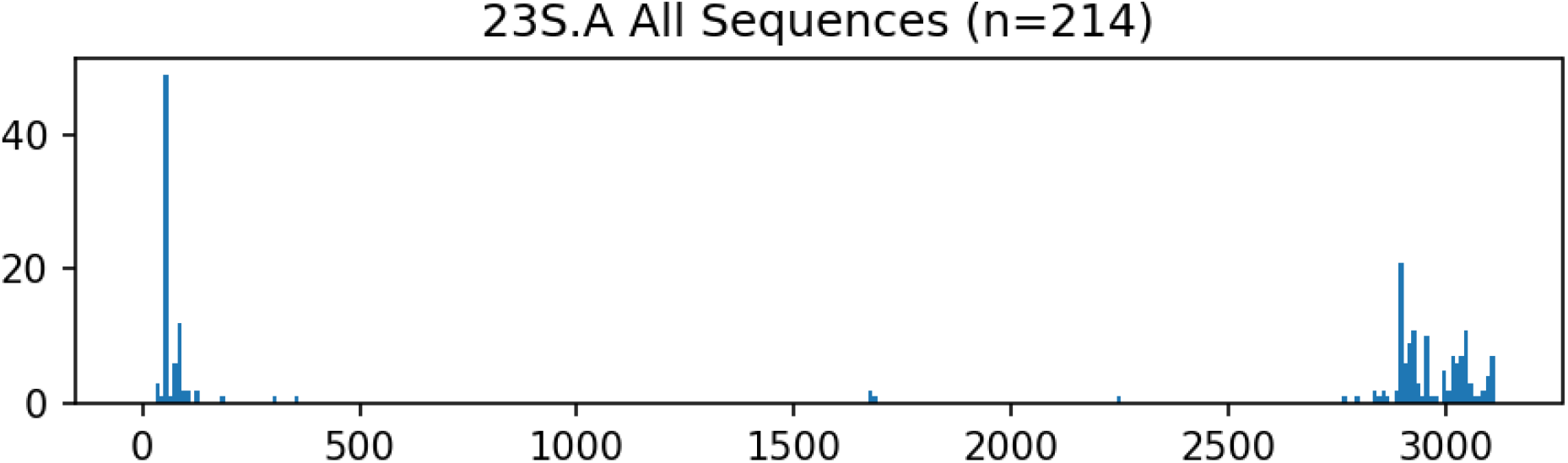
Sequence length histogram of the 23S.A dataset [2]

For the simulated datasets, we summarize the empirical properties in Supplementary Table 1. For the CRW biological datasets, we summarize the empirical properties in Supplementary Table 2.

#### MSA Methods

We compare our new MSA pipeline, HMMerge, to a collection of leading MSA methods. Specifically, we compare to: UPP, WITCH, MAGUS, PASTA, Clustal Omega [16], MUSCLE [4], MAFFT [6], and T-COFFEE [5]. The only methods that were not run in their default modes were MAFFT, which was run using the L-INS-i algorithm (to maximize accuracy), and T-COFFEE, which was run using the regressive mode. The exact commands and versions for each of the methods are supplied in the Supplementary Materials.

#### Computational Resources

All runs of HMMerge, WITCH, and UPP were run on the Illinois Campus Cluster with 16 cores available for parallelism and a default memory limit of 256GB. The standard methods (MAGUS, PASTA, MAFFT, Clustal Omega, T-COFFEE, and MUSCLE) were run on BlueWaters [1] for the ROSE and CRW datasets.

#### Evaluation Criteria

We evaluated the methods mainly on query sequence alignment error. Alignment error was calculated using FastSP [10]. FastSP is a tool that can calculate the SPFN and SPFP scores of an estimated alignment given a reference alignment, where SPFN and SPFP measure the proportion of false negative and false positive pair-wise homologies in the estimated alignment with respect to the reference alignment. Specifically, SPFN, or sum-of-pairs false negatives, is obtained by dividing the number of homologies found in the reference alignment but not in the estimated alignment by the number of homologies in the reference alignment, and SPFP, or sum-of-pairs false positives, is obtained by dividing the number of homologies found in the estimated alignment but not in the reference alignment by the homologies in the estimated alignment.

#### HMMerge and UPP Decomposition Parameters

For this study, we chose to do a centroid edge decomposition which is when the backbone tree is split at a centroid edge, an edge that divides the leaves into two as balanced as possible halves, recursively until a stopping criterion is met. In all experiments, the stopping criteria for HMMerge was when the subtree reached size 50 or smaller while for UPP it was when the smallest subtree reached size 2 or smaller. In addition, HMMerge used a disjoint decomposition that only kept the leaf partitions while UPP, as is default, used a hierarchical decomposition that keeps every level of partition in the decomposition. The decision to vary the decomposition parameters was due to the runtime constraints. For UPP, decomposing hierarchically down to size 2 has shown to improve alignment accuracy [14], and it may help the accuracy of HMMerge as well. However, the slow runtime of HMMerge made it not viable to hierarchically decompose down to the granularity of UPP since that would have created significantly more HMMs than keeping just the leaf HMMs. Therefore, we chose to stop the decomposition at size 50 and only keep the leaf HMMs.

## 4 Results

### 4.1 Experiment 1: Comparison of HMMerge, WITCH, and UPP for adding query sequences on simulated datasets with introduced fragmentation

#### 4.1.1 ROSE-HF Nucleotide Simulated Datasets

We observed that HMMerge was able to achieve a much lower SPFN error than UPP at a slight increase in SPFP error on ROSE high fragmentary simulated datasets across all model conditions as shown in Figure 3. WITCH trailed behind HMMerge, and UPP was the worst method out of the three methods tested.

**Figure 3:**
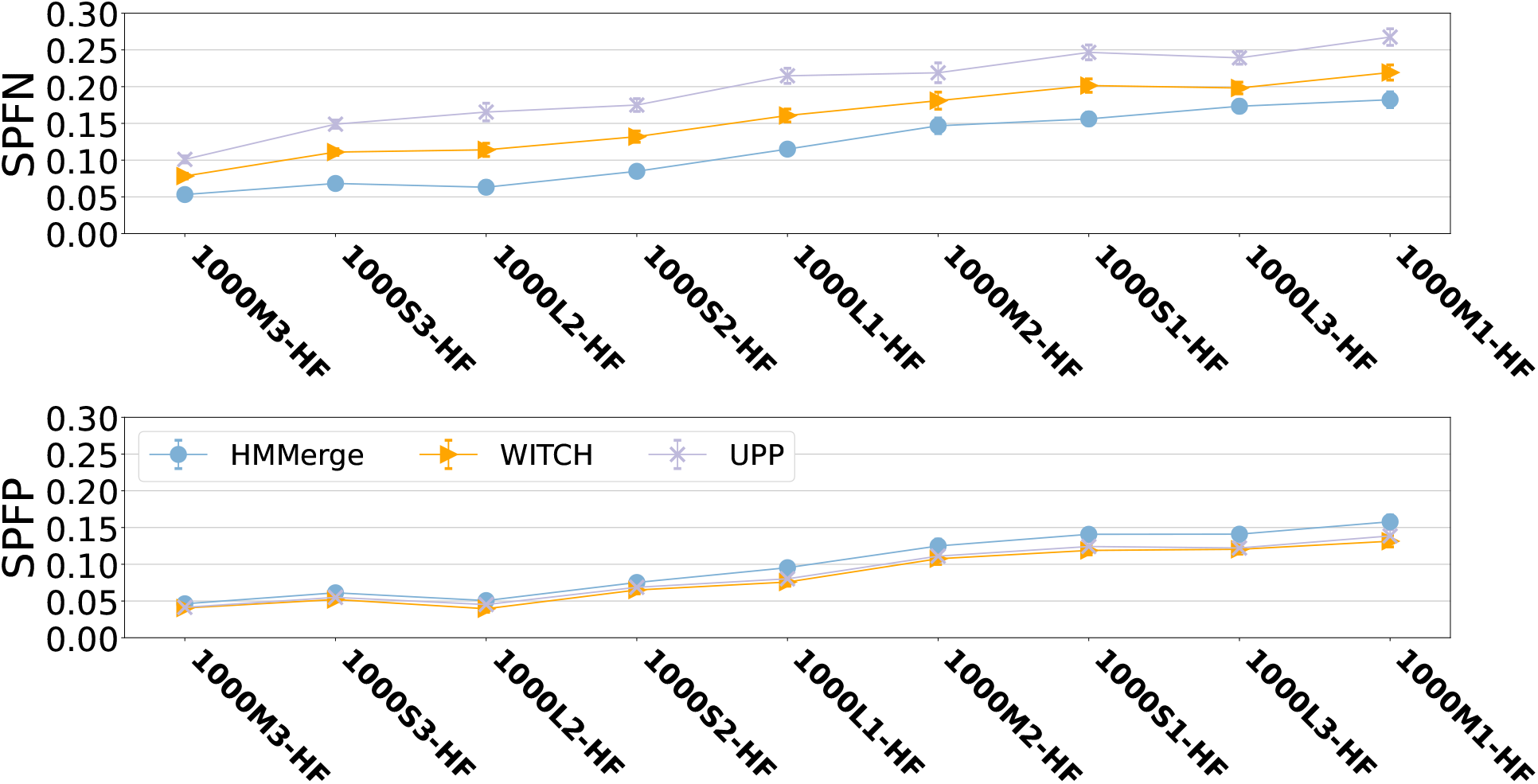
UPP, WITCH, and HMMerge query sequence alignment error on ROSE simulated datasets with introduced fragmentation. We show the average, SPFN, and SPFP error on the query sequences of the alignment produced by UPP, WITCH, and HMMerge on the ROSE HF simulated datasets.

#### 4.1.2 RNASim1000-HF and RNASim1000-UHF Datasets

On the RNASim1000-HF and RNASim1000-UHF datasets as shown in Table 3, there was a slight advantage to using HMMerge over WITCH and UPP in terms of SPFN error. HMMerge had the lowest average and SPFN error in the UHF condition. UPP had worse SPFN error and slightly better SPFP error compared to the other two methods and overall averaged to be competitive on the HF condition but worse on the UHF condition.

**Table 3:** Query sequence alignment error on RNASim1000 with introduced fragmentation. We show the average, SPFN, and SPFP error on the query sequences of the alignment produced by UPP, WITCH, and HMMerge on the RNASim1000-HF and RNASim1000-UHF simulated datasets.

|  | HMMerge | WITCH | UPP |
| --- | --- | --- | --- |
| RNASim1000-HF - average | **0.100** | **0.101** | **0.100** |
| RNASim1000-HF - SPFN | **0.102** | **0.103** | 0.108 |
| RNASim1000-HF - SPFP | 0.098 | 0.099 | **0.092** |
| RNASim1000-UHF - average | **0.103** | 0.105 | 0.108 |
| RNASim1000-UHF - SPFN | **0.107** | **0.108** | 0.123 |
| RNASim1000-UHF - SPFP | 0.099 | 0.102 | **0.093** |

### 4.2 Experiment 2: Comparison of HMMerge, WITCH, and UPP for adding query sequences on nucleotide biological datasets

#### 4.2.1 CRW Nucleotide Biological Datasets

We observed that on the CRW biological datasets (Table 4), HMMerge was slightly worse than WITCH and UPP on 23S.C, although the error rates of all methods were extremely close. However, HMMerge was much better than WITCH and UPP on 23S.A in terms of both average error and SPFN error. On 23S.A in particular, HMMerge was better than WITCH by all metrics - average, SPFN, and SPFP error and better than UPP in all but SPFP error. On the 5S.3, 5S.E, and 5S.T datasets, HMMerge in general did not perform as well as the other methods. HMMerge was, however, able to achieve lower SPFN error than UPP on all three 5S datasets. The full results on 5S.3, 5S.E, and 5S.T datasets are shown in the Supplementary Materials Section 4.3.

**Table 4:**
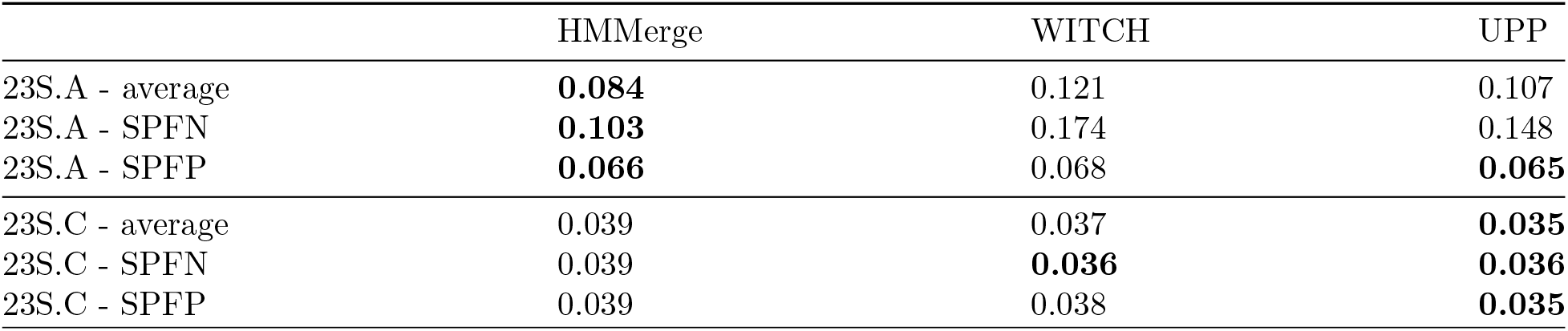
Query sequence alignment error on CRW biological datasets. We show the average, SPFN, and SPFP error on the query sequences of the alignment produced by UPP, WITCH, and HMMerge on the CRW biological datasets.

## 5 Discussion

The HMMerge pipeline is very closely related to UPP, differing from UPP only in how sequences are added into the backbone alignment.

A direct comparison among HMMerge, WITCH, and UPP is best made only with respect to the alignments they produce on the query sequences, since they are identical on the backbone sequences. On the ROSE high fragmentary simulated datasets, we see that HMMerge produces more accurate alignments than UPP or WITCH, with the biggest differences the conditions with high rates of evolution, which are also the conditions where alignment error tends to increase.

Overall, the strength of HMMerge in improving overall alignment error on the ROSE high fragmentary datasets comes from its ability to improve on SPFN scores, which can make up for the increased error on SPFP on some model conditions. In general, a lower SPFN score paired with a higher SPFP score relative to another method is a sign of over-alignment. Since HMMerge aims to align the entirety of the query sequence to a segment of the backbone alignment, a low SPFN score with a high SPFP score indicates that HMMerge is emitting a lot of characters through the match states rather than its insertion states whose emissions are considered non-homologous and hence do not count as true homologies for SPFN and SPFP score calculations.

On the RNASim1000-HF and RNASim1000-UHF datasets, we observed results in favor of HMMerge compared to WITCH and UPP, albeit at a lesser degree than the differences seen on the ROSE datasets. One possible explanation for this is that RNASim1000 datasets, although they have introduced fragmentation, was not simulated under fast evolving conditions such as those of ROSE and hence have lower p-distances.

It is worth noting that on the biological datasets tested, the results were mixed. One reason for the mixed results is that the biological datasets generally have relatively low average p-distances. For example, HMMerge was more effective on 23S.C than on 23S.A. We suspect that this is due to the lower average p-distance of 23S.A compared to 23S.C. In addition, not all biological datasets experience the kind of sequence length fragmentation such as the one that the simulated datasets underwent. This leads to an observation that HMMerge is most effective when used with datasets exhibiting high degree of fragmentation as well as high rates of evolution.

HMMerge is a method that can align query sequences which, when used in an UPP-like pipeline, shows great potential in improving the accuracy of alignments. On various simulated datasets we have tried, HM-Merge was able to achieve lower overall error than any of the standard methods we tried (see Supplementary Materials for more details). The improvement in alignment error was more apparent in the harder model conditions where the rate of evolution was high.

The ability of HMMerge to improve on alignment accuracy is especially noticeable when compared to UPP-add or WITCH-add. The main difference in this step is that HMMerge utilizes the information from all of the HMMs constructed from the backbone and uses the Viterbi algorithm whereas UPP-add and WITCH-add aligns extended alignments for each query sequence using *hmmalign*, which heuristically optimizes for the forward score in the forward-backward algorithm. This difference is potentially what gives HMMerge its edge.

One important theoretical difference between HMMerge and the UPP-add is that, although both methods rank the input HMMs, HMMerge ranks the HMMs through adjusted bit-scores, which accurately captures the probability of a query sequence being emitted by an HMM. UPP-add ranks the HMMs with the raw bit-scores, which UPP uses as a heuristic to assess the importance of an HMM.

## 6 Conclusion

HMMerge is a multiple sequence alignment method that can be used to add query sequences to backbone alignments. When used in a full pipeline, HMMerge can be used to align datasets with sequence length heterogeneity and achieve competitive, if not better, accuracy than current leading alignment methods, albeit often slower than other methods. HMMerge is especially effective under fast evolving and high fragmentary sequence length conditions where it excels in improving on SPFN scores, meaning that the true homologies represented by HMMerge are more reliable than other methods.

Although we used a subset set size of 50 for the decomposition and used a centroid-edge disjoint decomposition, meaning that we recursively split at a centroid edge and only kept the leaves of the subsets, the HMMerge algorithm naturally works with any kind of decomposition. Namely, it is able to handle arbitrary sized subsets that do not have to be disjoint and can be hierarchical as is the default in UPP. It is possible that varying the decomposition size and type could lead to runtime improvements and even further accuracy improvements.

## Supporting information

are shown in the Supplementary Materials

## 8 Author contributions statement

M.P. and T.W. conceived of HMMerge, M.P. implemented the algorithm and conducted the experiments, M.P. and T.W. wrote and reviewed the manuscript.

## 9 Acknowledgments

This work was supported by a subaward from Sandia National Labs (LDRD to Kelly Williams) and the US National Science Foundation [grant number 2006069].

