## Supplementary material for "HMMerge: an Ensemble Method for Improving Multiple Sequence Alignment": are shown in the Supplementary Materials

#### Contents

**1 Commands and Versions of Software Used** **3**

**2 Edge Re-normalization for $p_{entry}$ and $p_{exit}$** **5**

**3 Additional Tables** **6**

**4 Additional Figures** **7**

### 1 Commands and Versions of Software Used

**UPP** Version: 4.5.2
Availability: https://github.com/smirarab/sepp

```
python run_upp.py -c <Config file>
```

```
[commandline]
tree=<FastTree backbone tree>
alignment=<MAGUS backbone alignment>
sequence_file=<Unaligned query sequence file>
backboneSize=<backbone size>
alignmentSize=2
molecule=dna
cpu=16
outdir=<Output directory>
tempdir=<Temporary output directory>
```

**MAGUS** Version: 0.1.0b2
Availability: https://github.com/vlasmirnov/MAGUS

```
python <git root>/magus.py -d <Output directory> -i \
<Unaligned sequences> -o <Output filename>
```

**PASTA** Version: 1.9.0
Availability: https://github.com/smirarab/pasta

```
python run_pasta.py -i <Unaligned sequences> --num-cpus 16 \
-o <Output directory> --temporaries <Temporary output directory>
```

**MAFFT** Version: 7.487
Availability: https://mafft.cbrc.jp/alignment/software/linux.html

```
linsi --thread 16 <Unaligned sequences> 1> <Output filename>
```

**Clustal Omega** Version: 1.2.4
Availability: http://www.clustal.org/omega/#Download

```
clustalo --threads=16 --in <Unaligned sequences> --out \
<Output filenanme>
```

**T-COFFEE** Version: 13.45.0.4846264
Availability: https://www.tcoffee.org/Packages/Stable/Latest/

```
t_coffee -thread=16 -reg -seq <Unaligned sequences> -outfile \
<Output filename>
```

**MUSCLE** Version: 3.8.31
Availability: https://drive5.com/muscle/downloads_v3.htm

```
muscle -in <Unaligned sequences> -out <Output filename>
```

**FastSP** Version: 1.7.1
Availability: https://github.com/smirarab/FastSP
Note: When evaluating MAFFT estimated alignments, the ml and mlr flags should be omitted.

```
java -jar FastSP.jar -r <Reference alignment> -e \
<Estimated alignment> -ml -mlr
```

**HMMerge** Commit ID: 0457b96a63388ebbb91cfe270e6c135a3bd5ff4c
Availability: https://github.com/MinhyukPark/HMMerge

```
python <git root>/main.py --input-dir <Directory with \
partitioned backbone alignments> --backbone-alignment \
<Backbone alignment> --query-sequence-file \
<Query sequences> --output-prefix <Output directory> \
--num-processes 16 --model DNA
```

**WITCH** Commit ID: a5fff8b8e9491869a151318061543a2af22db7df
Availability: https://github.com/c5shen/WITCH

```
python <git root>witch.py -t 16 -b <Backbone alignment> \
-e <Backbone tree> -q <Query sequences> -d \
<Output directory> --molecule dna -o <Output filename>
```

#### 2 Edge Re-normalization for $p_{entry}$ and $p_{exit}$

Let $T_0$ be the old transition probability matrix, $T_1$ be the new transition probability matrix, $n$ is the number of states as stated before, and $S$ be the number of match states.

$$T_1[0, i] = \begin{cases} p_{entry} & i \text{ is a match state,} \\ (1 - S \cdot p_{entry}) \cdot \frac{T_0[0,i]}{\Sigma T_0[0,\cdot]} & i \text{ is not a match state} \end{cases}$$

For the exit probabilities, we do the following modification in order to re-normalize old edge weight such that outgoing egde weights sum to 1.

$$T_1[i, j] = \begin{cases} p_{exit} & i \text{ is not an insertion state} \\ & \text{and } j = n, \\ (1 - p_{exit}) \cdot \frac{T_0[i,j]}{\Sigma T_0[i,\cdot]} & i \text{ is not an insertion state} \\ & \text{and } j \neq n \end{cases}$$

#### 3 Additional Tables

Table 1: **Simulated DNA/RNA Dataset Overview** Here, we show the basic empirical statistics about the datasets used in this study. P-distances refer to the normalized Hamming distance such that two identical sequences have a p-distance of 0 while two sequences with no shared characters have a p-distance of 1. The p-distances shown are prior to fragmentation.

| Name | # Sequences | # Full | # Query | avg. p-dist. | max. p-dist. | avg len. | median len. | stddev len. |
|---|---|---|---|---|---|---|---|---|
| 1000S1 | 1000 | 500 | 500 | 0.694 | 0.768 | 1002 | 1002 | 5 |
| 1000S2 | 1000 | 500 | 500 | 0.693 | 0.768 | 1002 | 1001 | 4 |
| 1000S3 | 1000 | 500 | 500 | 0.686 | 0.763 | 1002 | 1002 | 4 |
| 1000S4 | 1000 | 500 | 500 | 0.501 | 0.608 | 1001 | 1000 | 2 |
| 1000S5 | 1000 | 500 | 500 | 0.498 | 0.611 | 1001 | 1000 | 2 |
| 1000M1 | 1000 | 500 | 500 | 0.695 | 0.769 | 1011 | 1009 | 12 |
| 1000M2 | 1000 | 500 | 500 | 0.684 | 0.762 | 1014 | 1012 | 14 |
| 1000M3 | 1000 | 500 | 500 | 0.660 | 0.741 | 1008 | 1005 | 10 |
| 1000M4 | 1000 | 500 | 500 | 0.495 | 0.606 | 1007 | 1005 | 9 |
| 1000M5 | 1000 | 500 | 500 | 0.499 | 0.602 | 1004 | 1002 | 7 |
| 1000L1 | 1000 | 500 | 500 | 0.695 | 0.769 | 1015 | 1015 | 17 |
| 1000L2 | 1000 | 500 | 500 | 0.696 | 0.769 | 1007 | 1003 | 11 |
| 1000L3 | 1000 | 500 | 500 | 0.687 | 0.763 | 1032 | 1027 | 26 |
| 1000L4 | 1000 | 500 | 500 | 0.500 | 0.608 | 1008 | 1004 | 12 |
| 1000L5 | 1000 | 500 | 500 | 0.496 | 0.606 | 1006 | 1000 | 11 |
| RNASim1000 | 1000 | 500 | 500 | 0.411 | 0.609 | 1554 | 1554 | 10 |

Table 2: **Biological DNA/RNA Dataset Overview** Here, we show the basic empirical statistics about the datasets used in this study. P-distances refer to the normalized Hamming distance such that two identical sequences have a p-distance of 0 while two sequences with no shared characters have a p-distance of 1. The p-distances shown are prior to fragmentation. For these biological datasets, the backbone sequences were chosen by partitioning the datasets at length 1250 for 23S.C and 23S.A and at length 100 for 5S.3, 5S.E, and 5S.T.

| Name | # Sequences | # Full | # Query | avg. p-dist. | max. p-dist. | avg len. | median len. | stddev len. |
|---|---|---|---|---|---|---|---|---|
| 23S.A | 214 | 133 | 81 | 0.293 | 0.667 | 1851 | 2897 | 1397 |
| 23S.C | 374 | 274 | 100 | 0.143 | 0.750 | 2087 | 2642 | 1106 |
| 5S.3 | 5507 | 4439 | 1068 | 0.417 | 1.000 | 106 | 118 | 31 |
| 5S.E | 2774 | 1886 | 888 | 0.305 | 1.000 | 96 | 119 | 39 |
| 5S.T | 5751 | 4677 | 1074 | 0.425 | 1.000 | 106 | 119 | 31 |

### 4 Additional Figures

#### 4.1 ROSE datasets with introduced fragmentation

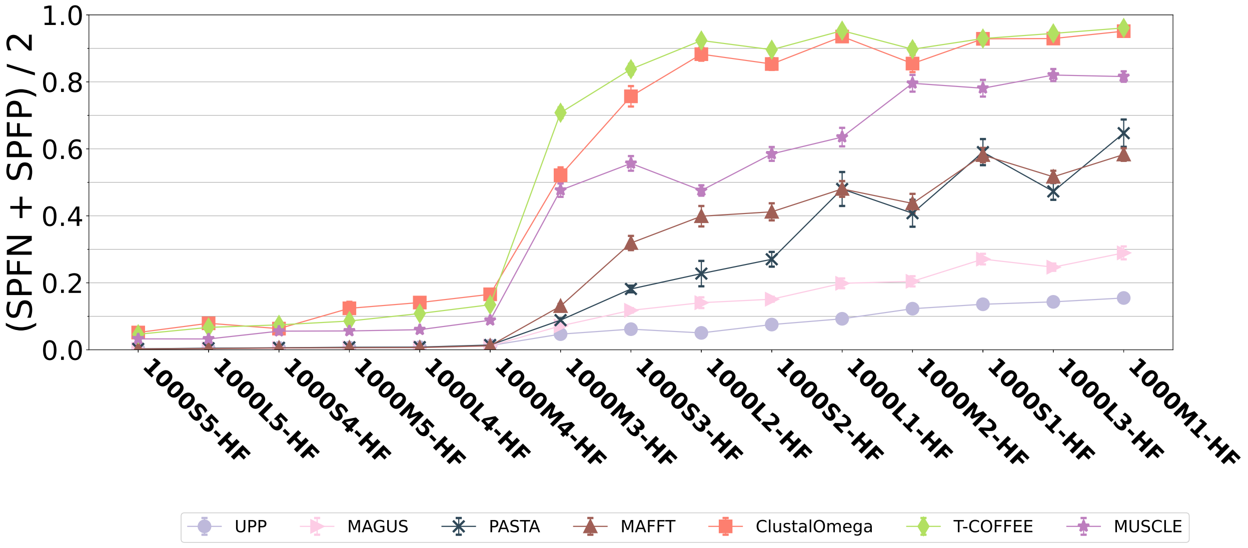

Figure 1: **Alignment error for benchmark methods on ROSE simulated datasets with introduced fragmentation** Alignment error (average of SPFN and SPFP) of the final alignment produced by each method. These are highly fragmented ("HF" for short) datasets created from ROSE simulated datasets introduced in the SATé study [1]. All datasets have 1000 sequences, with rates of evolution that increase from left-to-right. The error bars indicate standard error over 20 replicates.

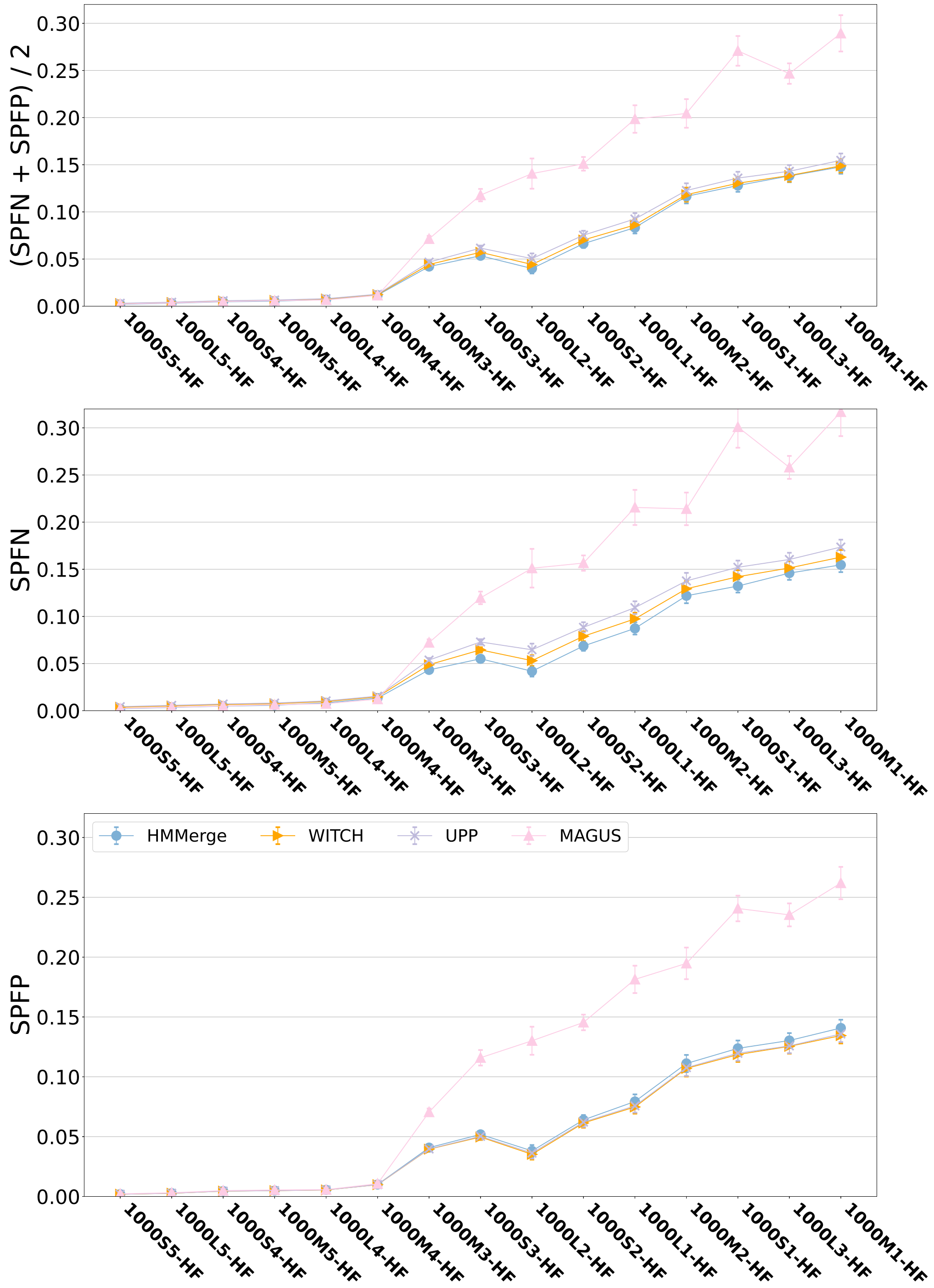

Figure 2: **Total alignment error for HMMerge, WITCH, UPP, and MAGUS on ROSE simulated datasets with introduced fragmentation** We show the alignment error on the final alignment produced by each of the top four methods (HMMerge, WITCH, UPP, and MAGUS). The error bars indicate standard error over 20 replicates.

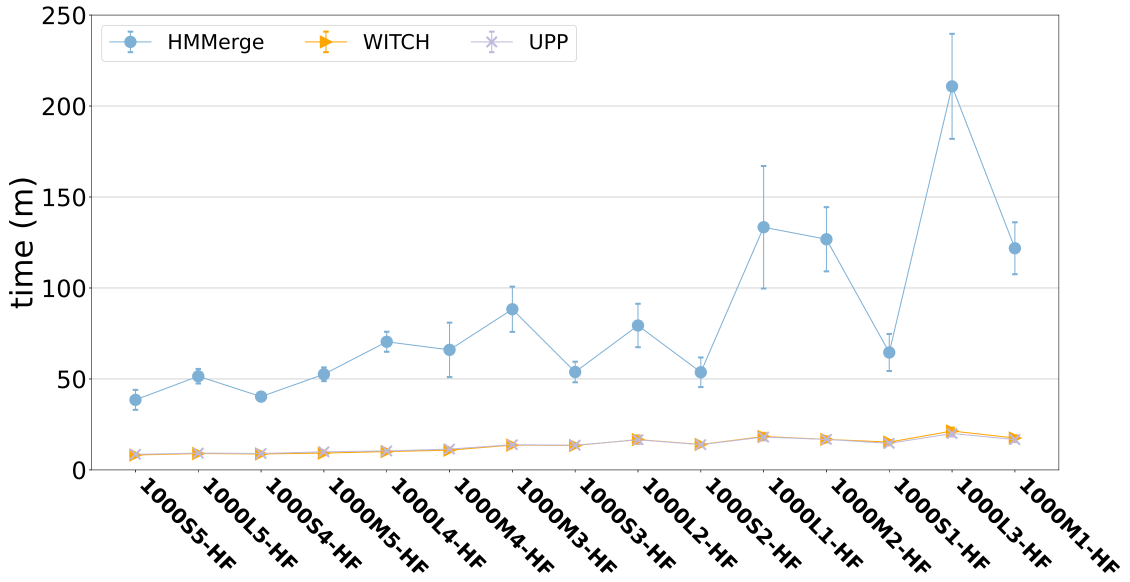

Figure 3: **Total Runtime for HMMerge, WITCH, and UPP pipelines on ROSE simulated datasets with introduced fragmentation** We show the runtime of obtaining the final alignment by HMMerge, WITCH, and UPP; this includes the time required for building the backbone alignment with MAGUS and tree with FastTree. The error bars indicate standard deviation over 20 replicates.

#### 4.2 RNASim1000 datasets with introduced fragmentation

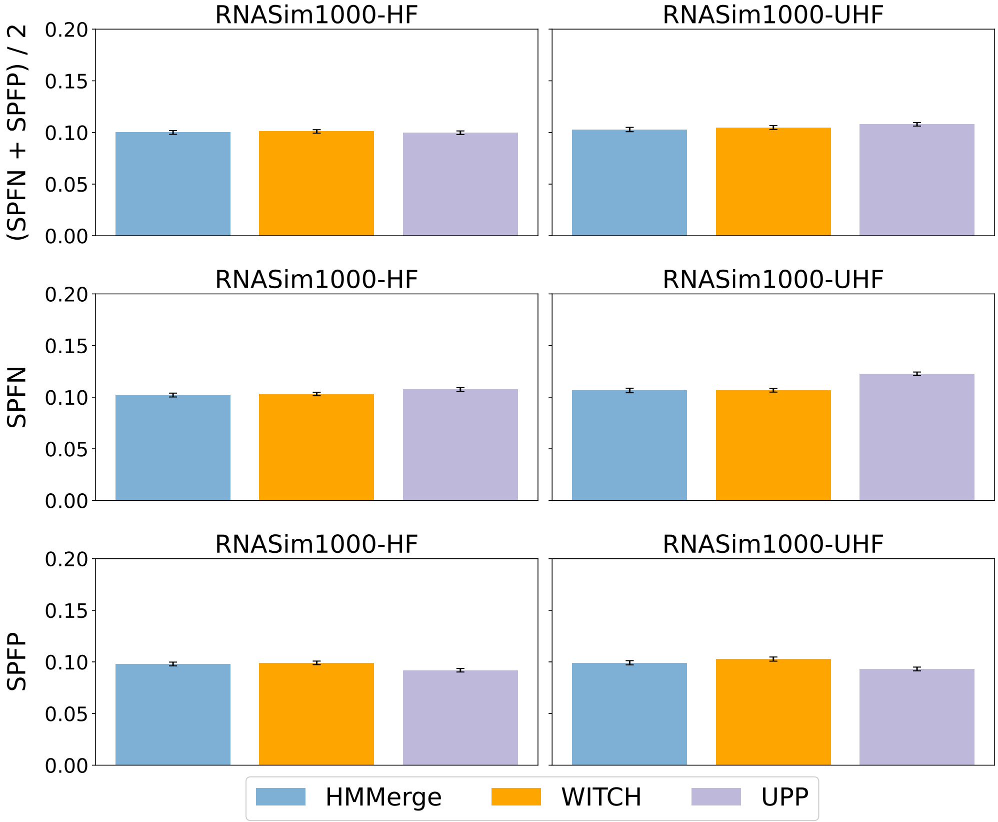

Figure 4: **UPP, WITCH, and HMMerge query sequence alignment error on RNASim1000 simulated datasets with introduced fragmentation** We show the average, SPFN, and SPFP error on the query sequences of the alignment produced by UPP, WITCH, and HMMerge on the RNASim1000-HF and RNASim1000-UHF simulated datasets.

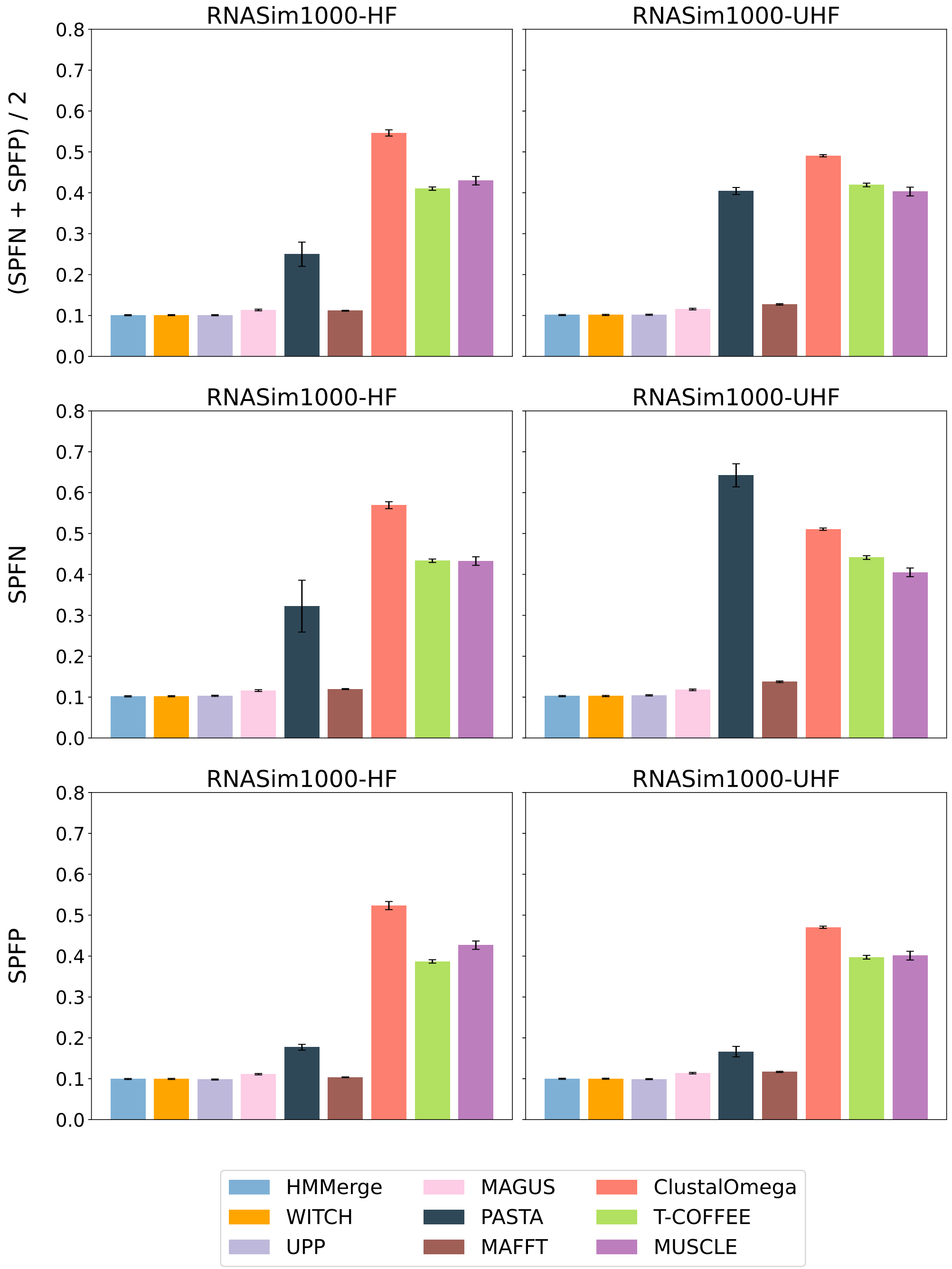

Figure 5: **Total alignment error for HMMerge, WITCH, UPP, and MAGUS on RNASim1000 simulated datasets with introduced fragmentation** We show the alignment error on the final alignment produced by each of the top four methods (HMMerge, WITCH, UPP, and MAGUS). The error bars indicate standard error over 10 replicates.

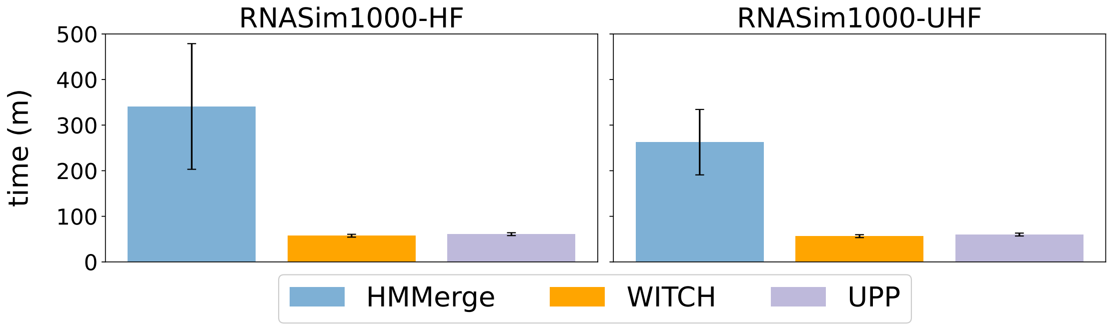

Figure 6: **Total Runtime for HMMerge, WITCH, and UPP pipelines on RNASim1000 simulated datasets with introduced fragmentation** We show the runtime of obtaining the final alignment by HMMerge, WITCH, and UPP; this includes the time required for building the backbone alignment with MAGUS and tree with FastTree. The error bars indicate standard deviation over 10 replicates.

##### 4.3 CRW datasets without introduced fragmentation

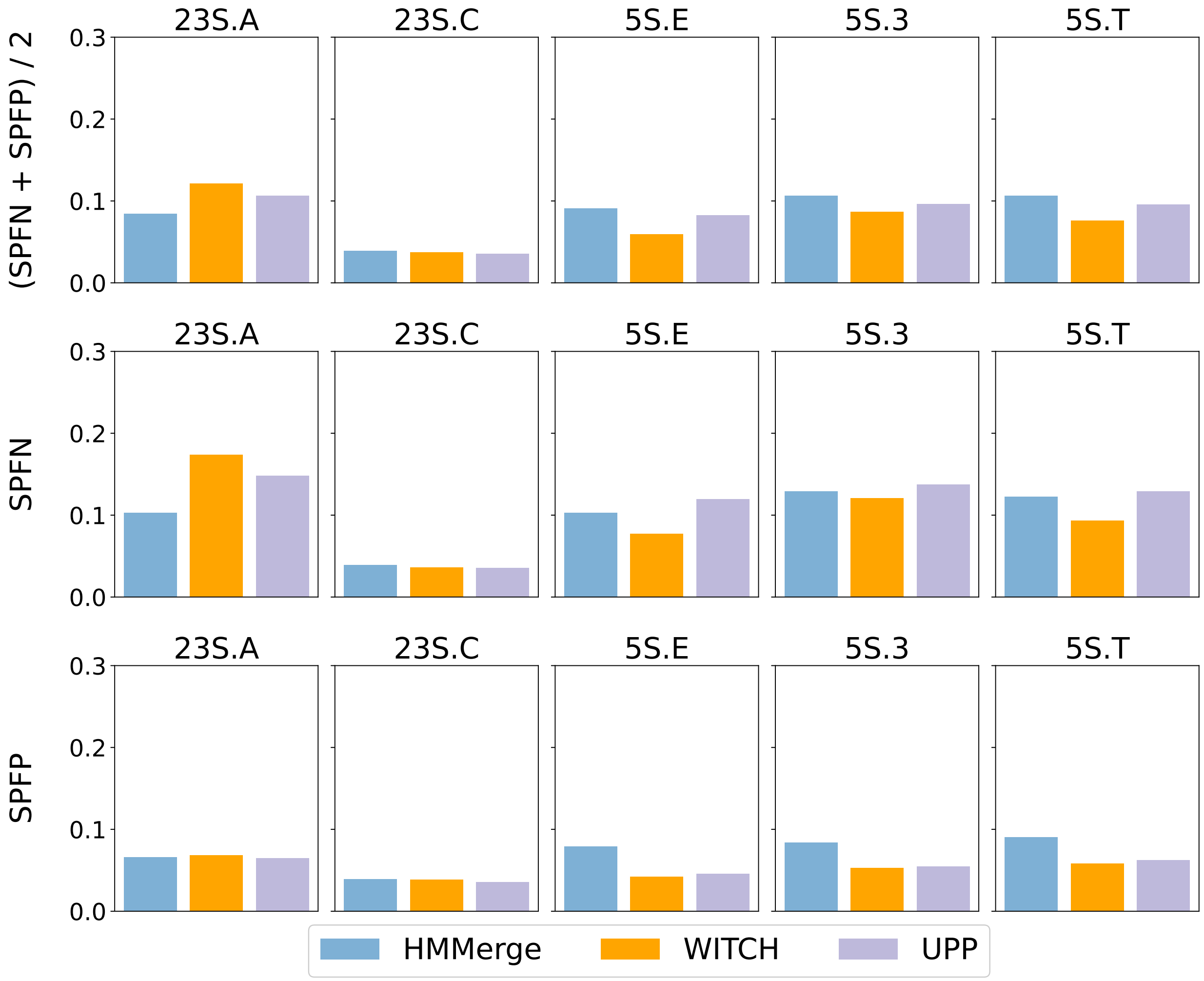

Figure 7: **UPP, WITCH, and HMMerge query sequence alignment error on CRW biological datasets with no introduced fragmentation** We show the average, SPFN, and SPFP error on the query sequences of the alignment produced by UPP, WITCH, and HMMerge on the CRW biological datasets with no introduced fragmentation.

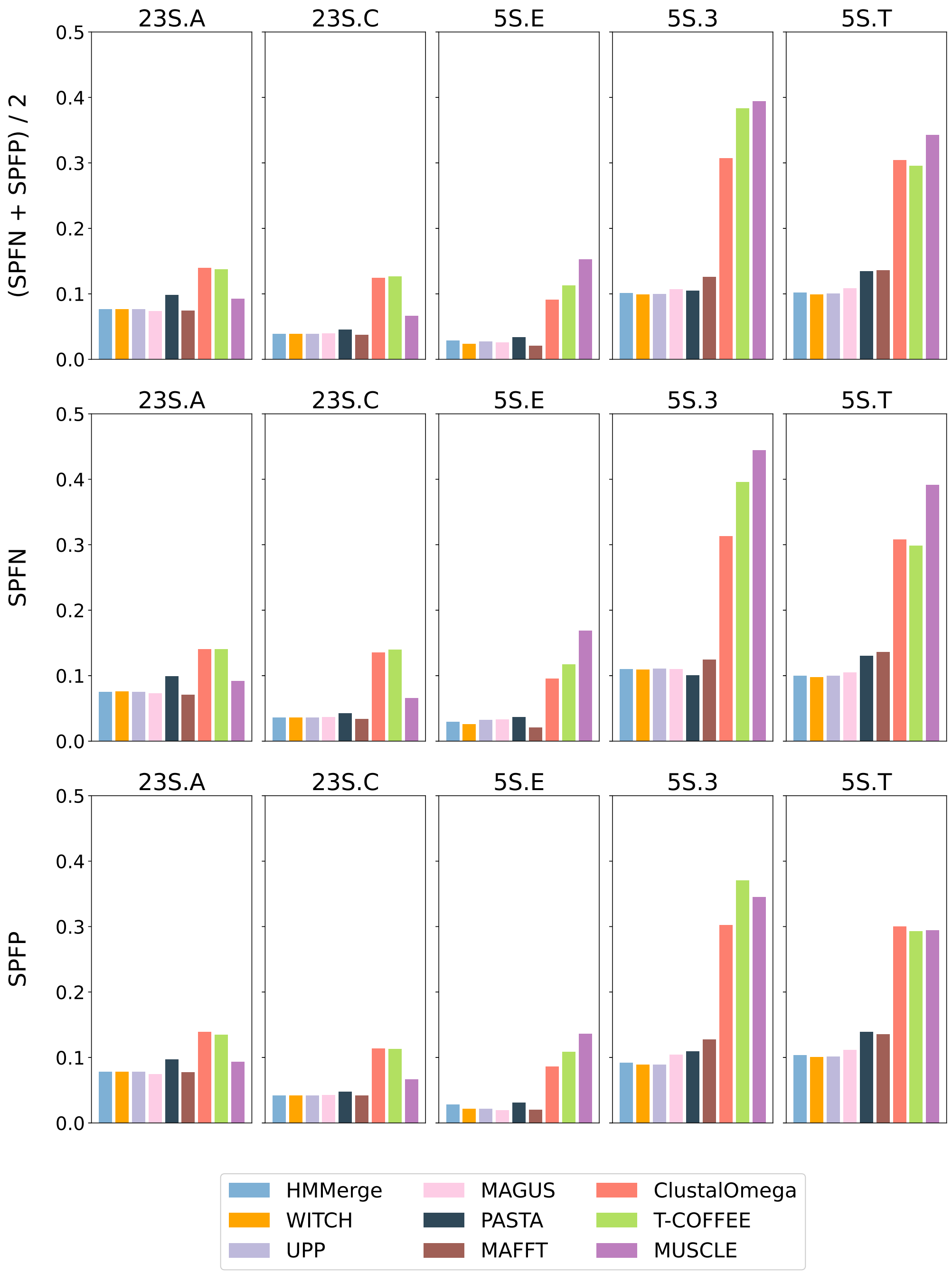

Figure 8: **Total alignment error on CRW biological datasets with no introduced fragmentation** We show the alignment error on the final alignment produced by each of the methods.

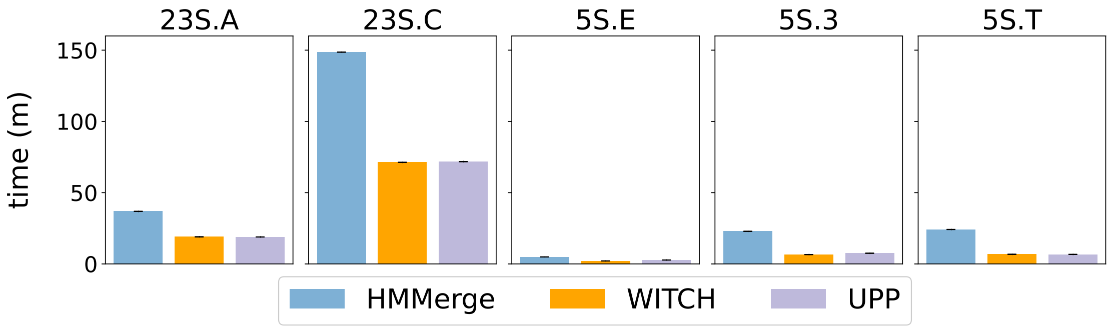

Figure 9: **Total runtime for HMMerge, WITCH, and UPP pipelines on CRW biological datasets with no introduced fragmentation** We show the runtime of obtaining the final alignment by HMMerge, WITCH, and UPP; this includes the time required for building the backbone alignment with MAGUS and tree with FastTree.

#### 4.4 ROSE Simulated Datasets Sequence Length Histograms

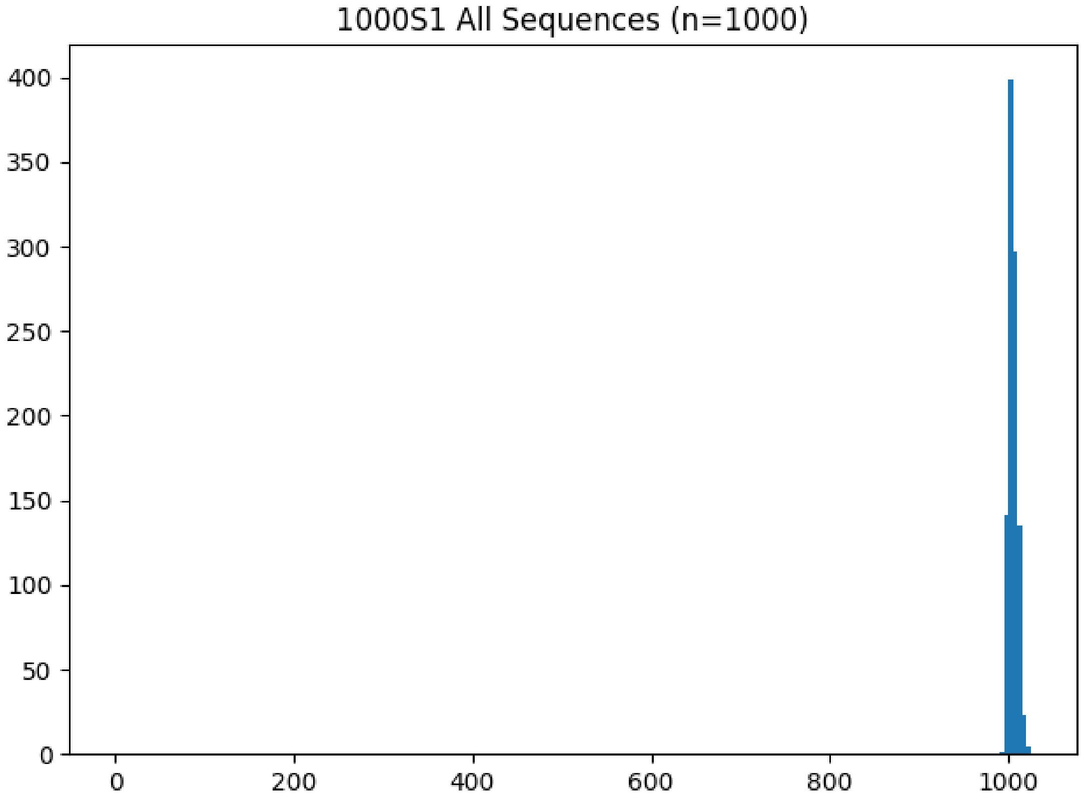

Figure 10: **ROSE 1000S1 histogram**

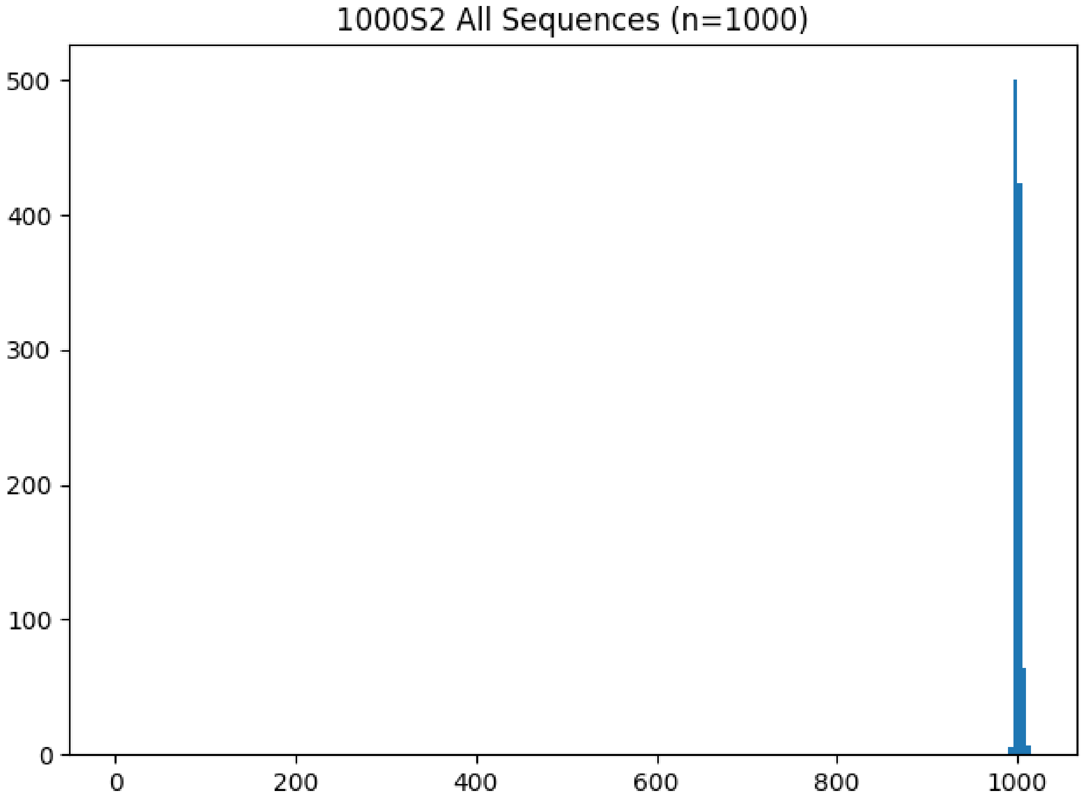

Figure 11: **ROSE 1000S2 histogram**

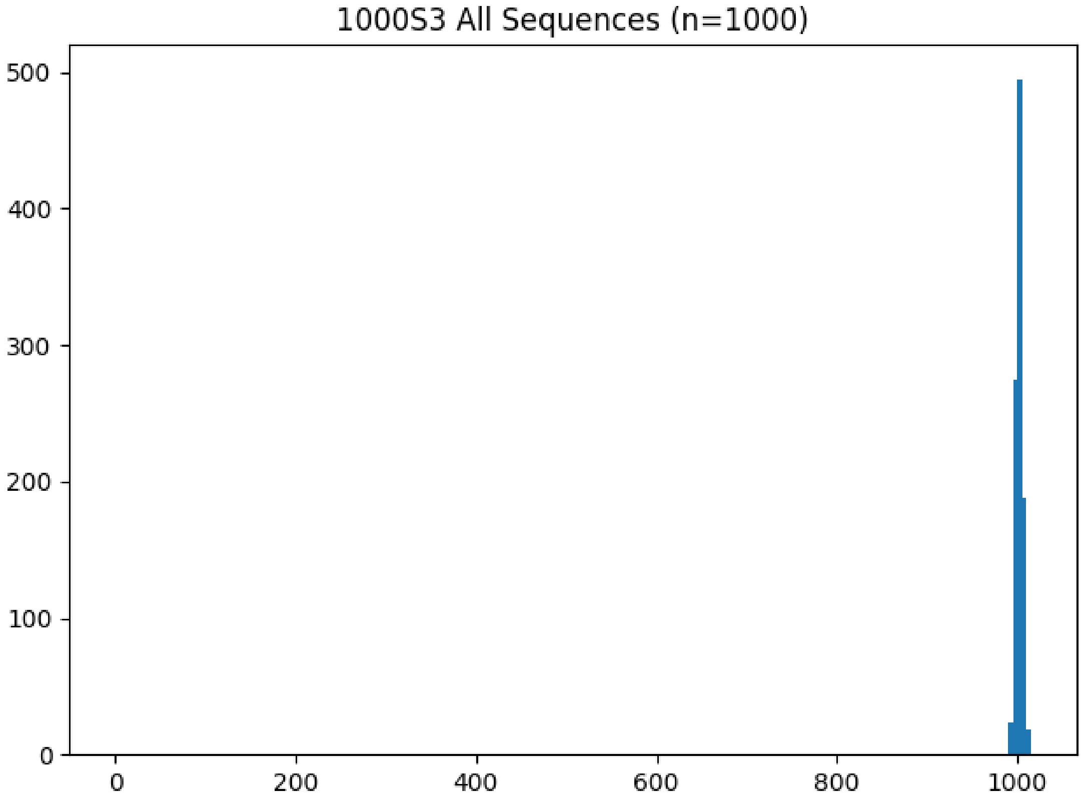

Figure 12: **ROSE 1000S3 histogram**

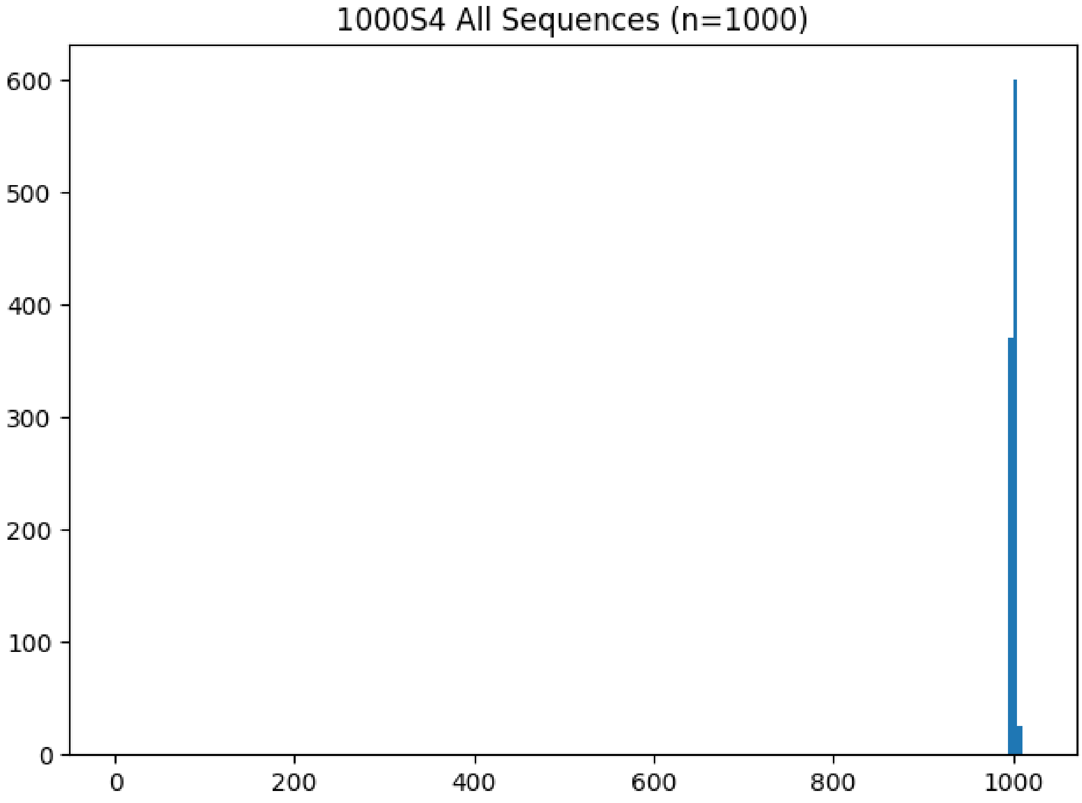

Figure 13: **ROSE 1000S4 histogram**

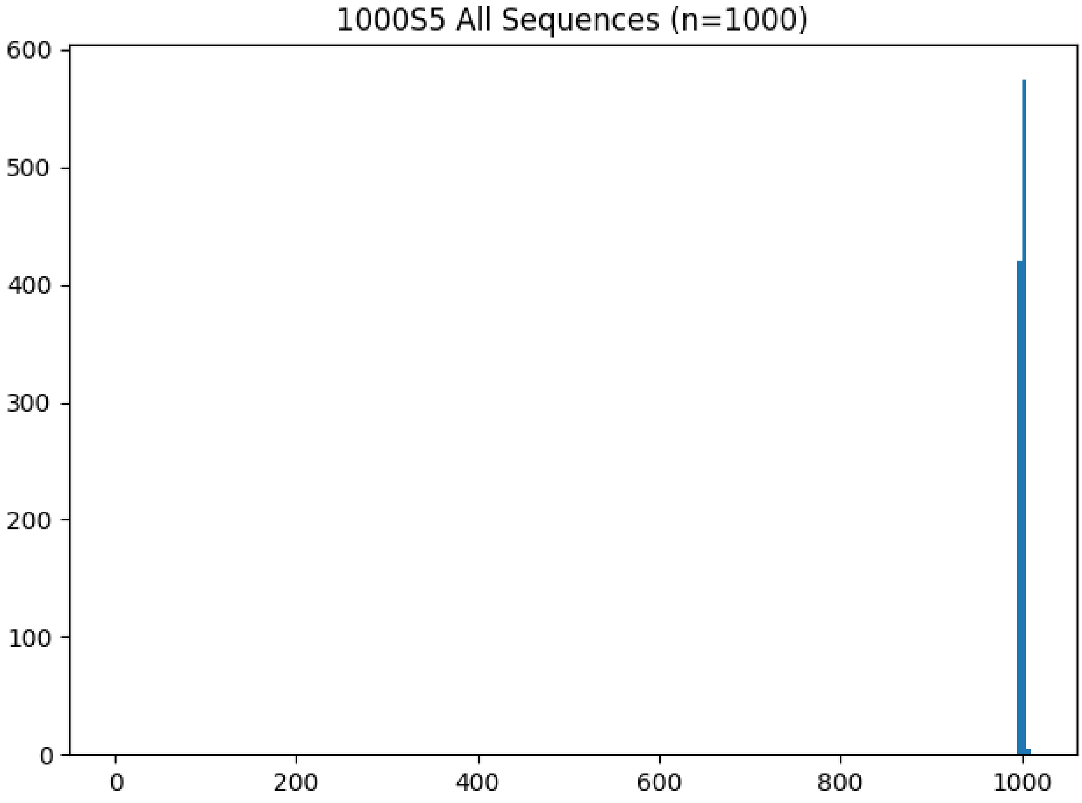

Figure 14: **ROSE 1000S5 histogram**

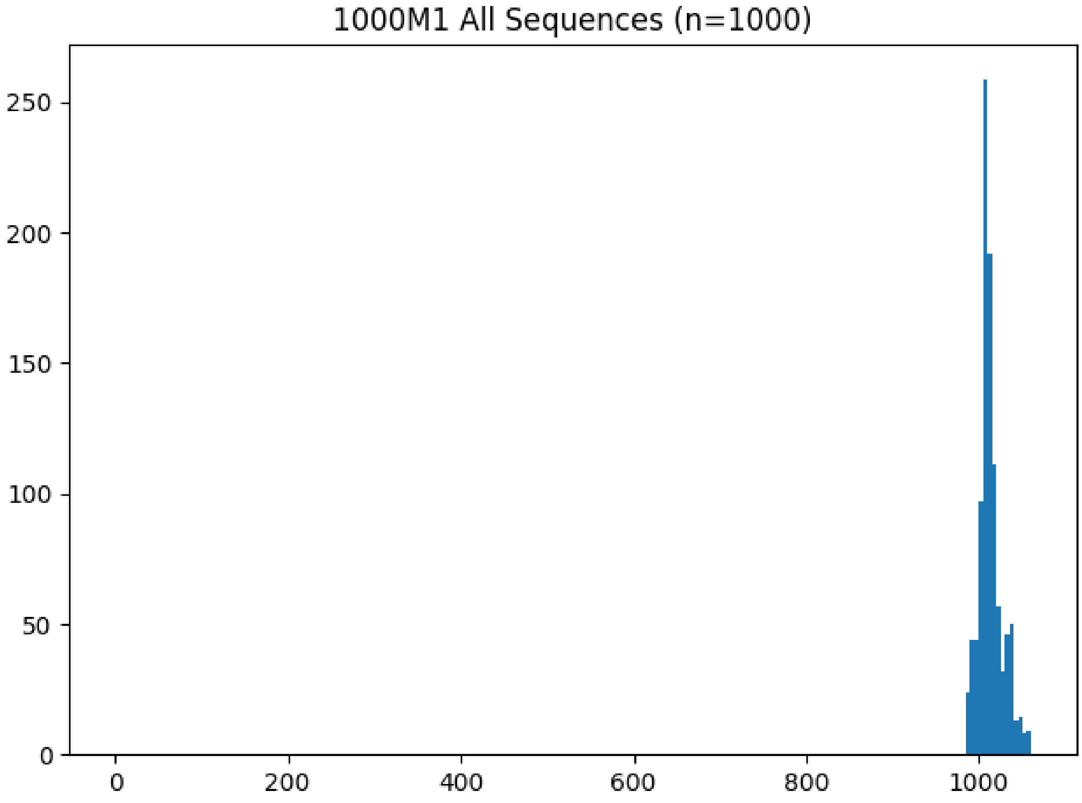

Figure 15: **ROSE 1000M1 histogram**

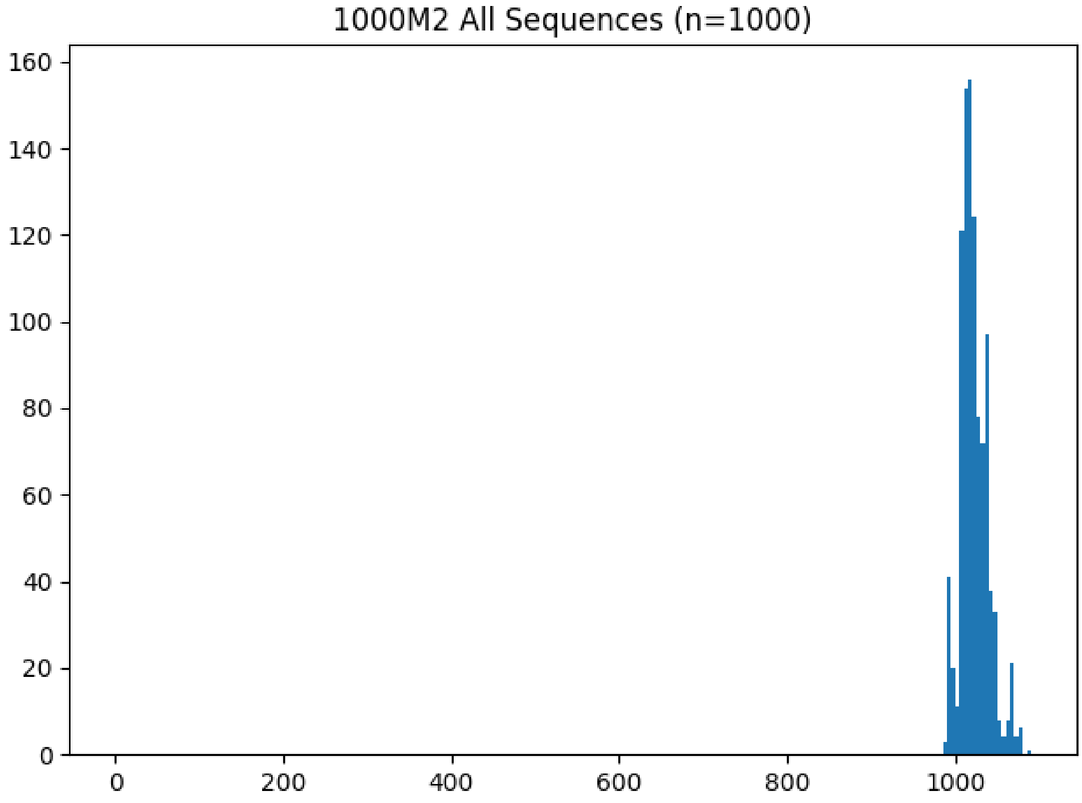

Figure 16: **ROSE 1000M2 histogram**

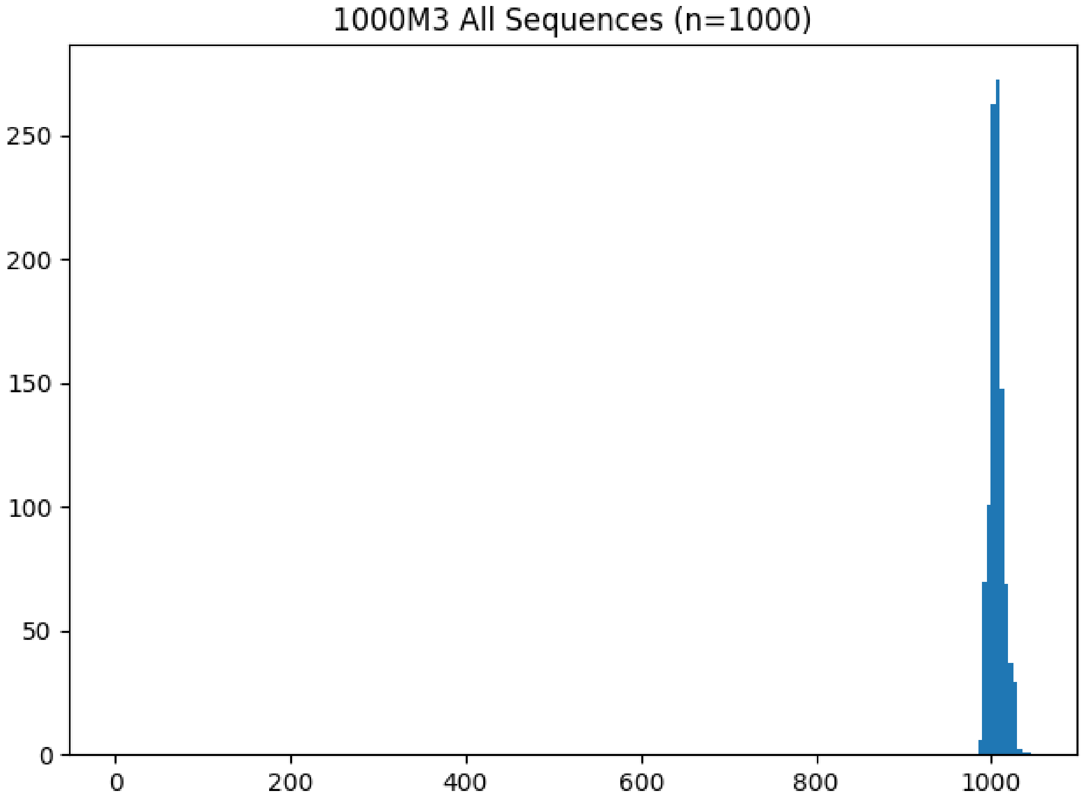

Figure 17: **ROSE 1000M3 histogram**

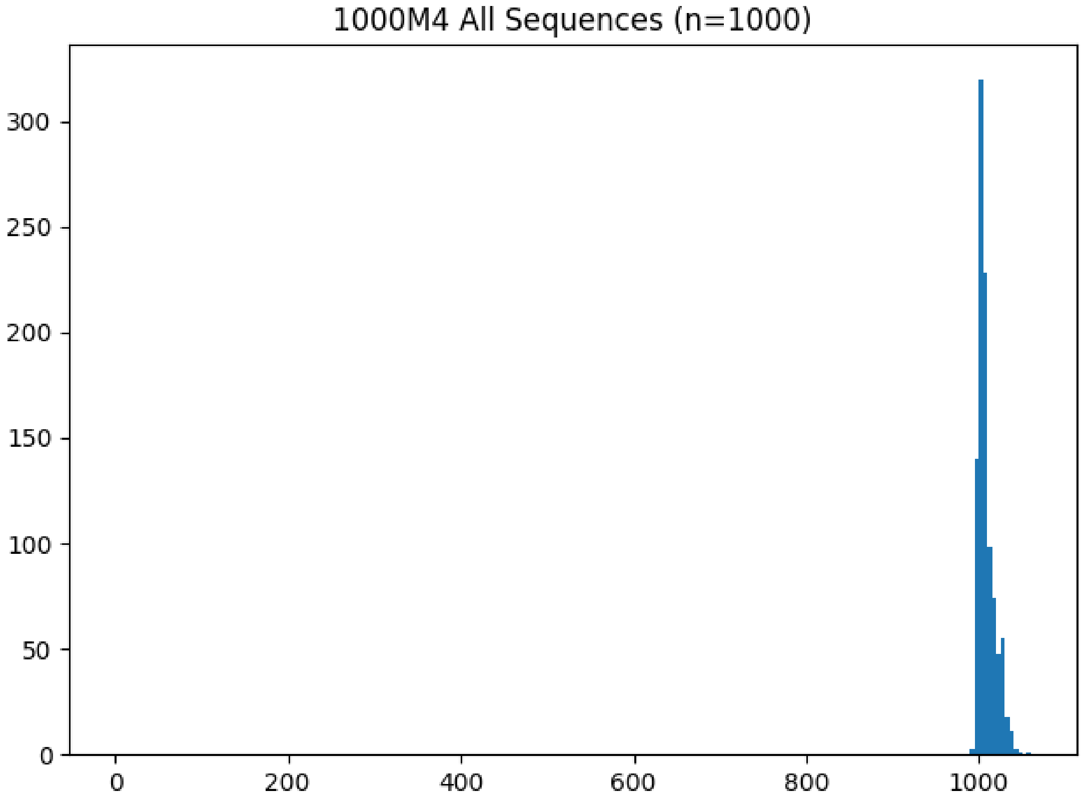

Figure 18: **ROSE 1000M4 histogram**

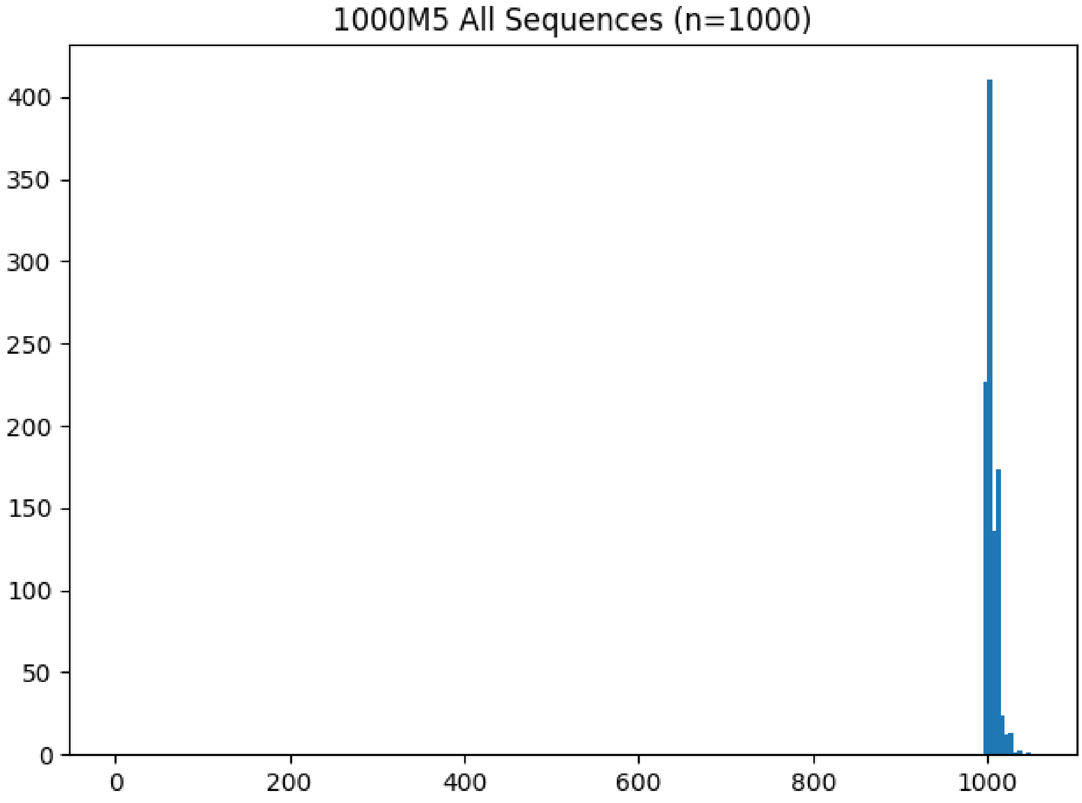

Figure 19: **ROSE 1000M5 histogram**

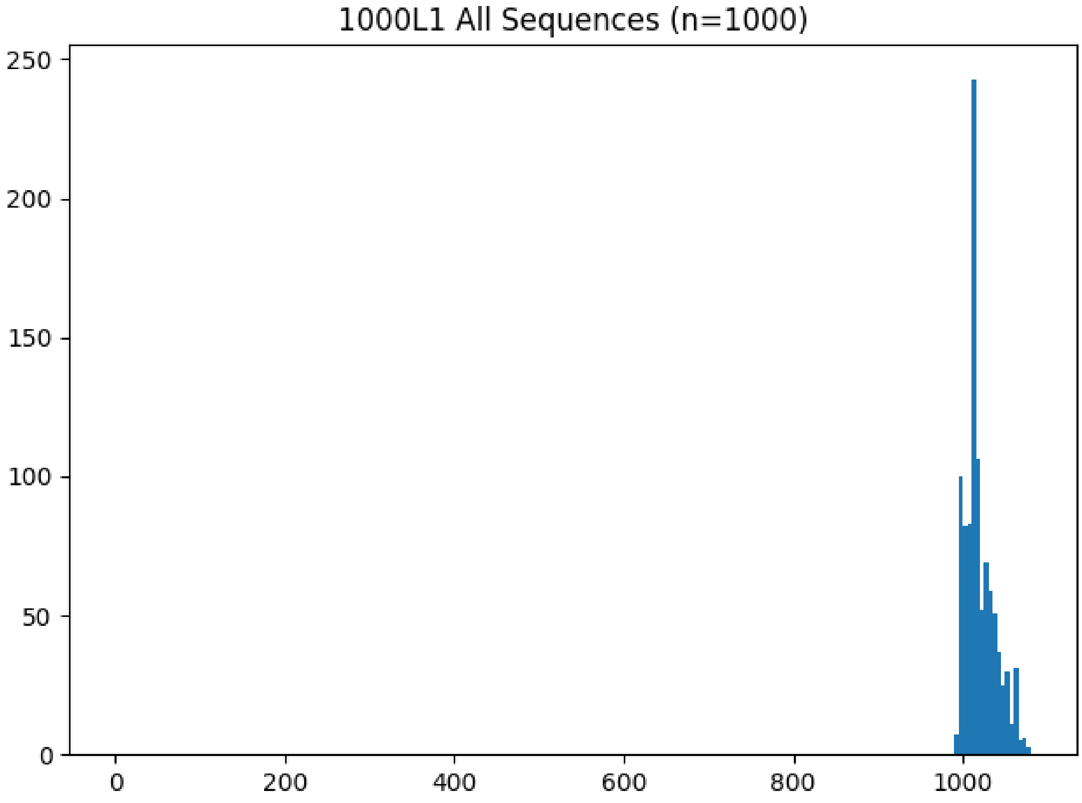

Figure 20: **ROSE 1000L1 histogram**

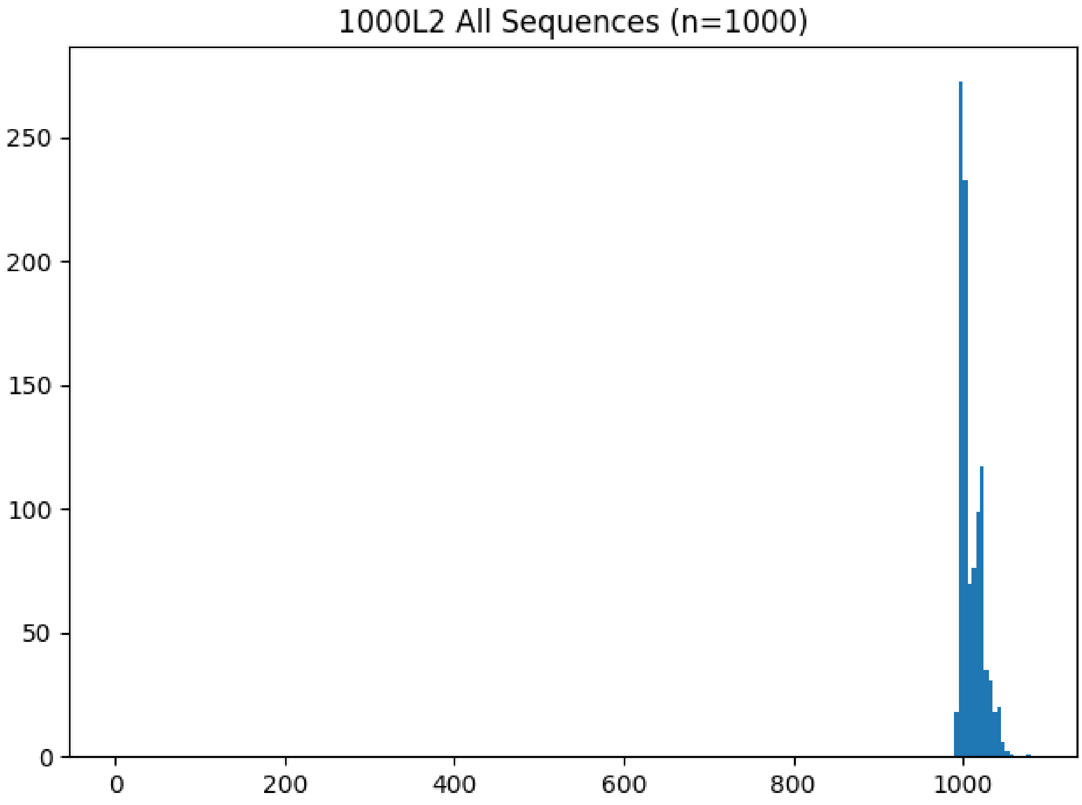

Figure 21: **ROSE 1000L2 histogram**

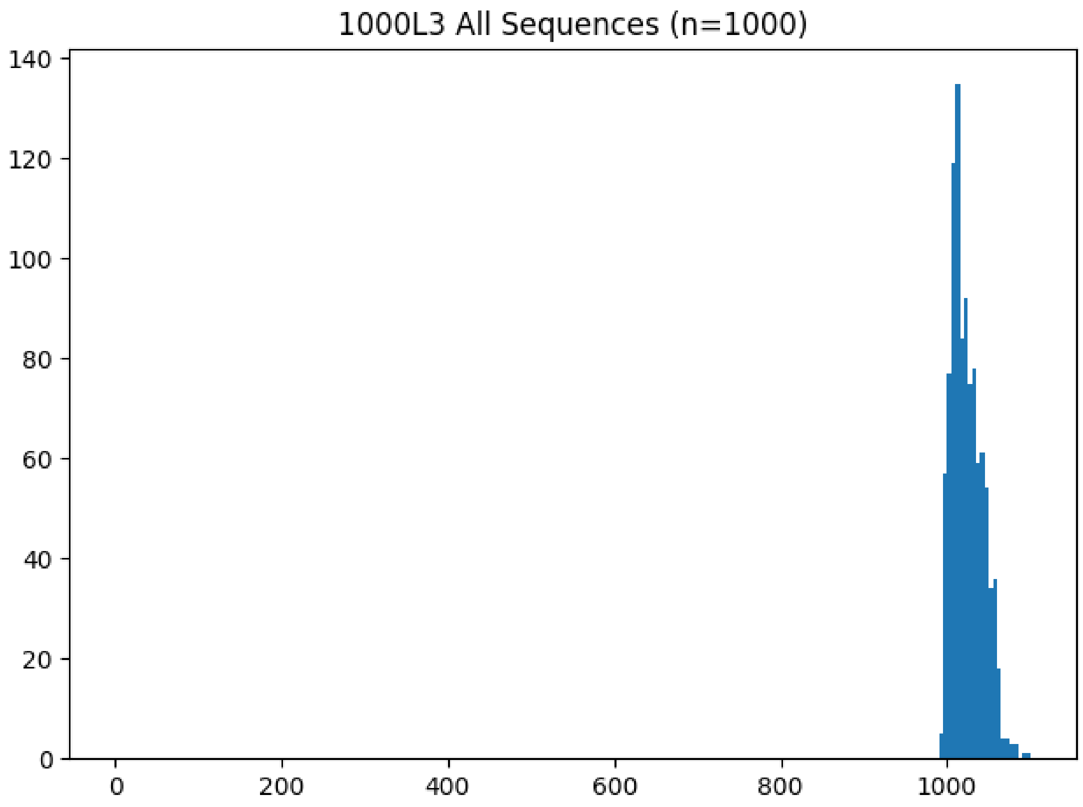

Figure 22: **ROSE 1000L3 histogram**

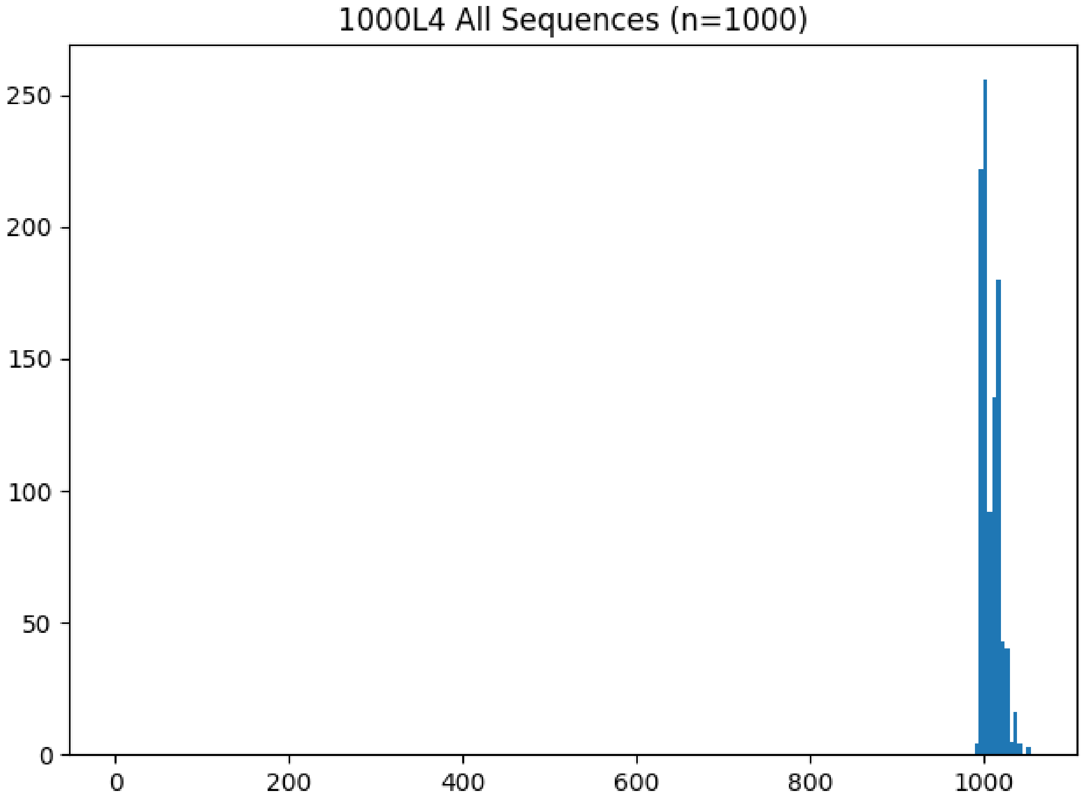

Figure 23: **ROSE 1000L4 histogram**

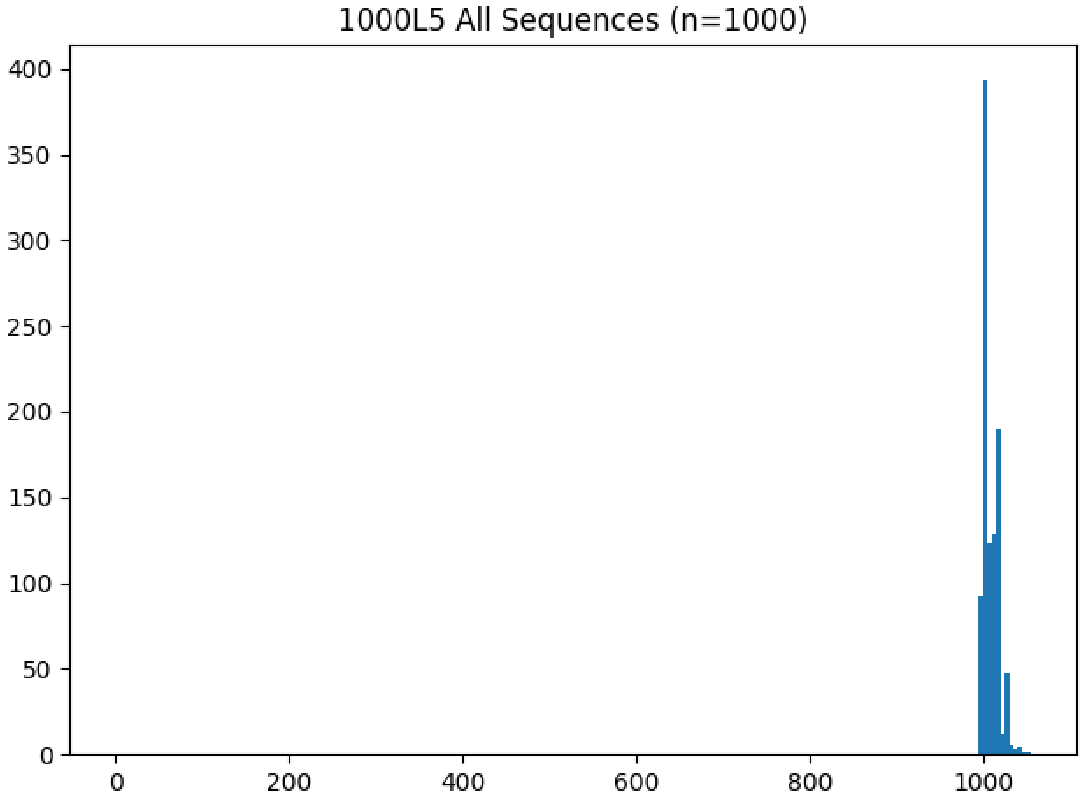

Figure 24: **ROSE 1000L5 histogram**

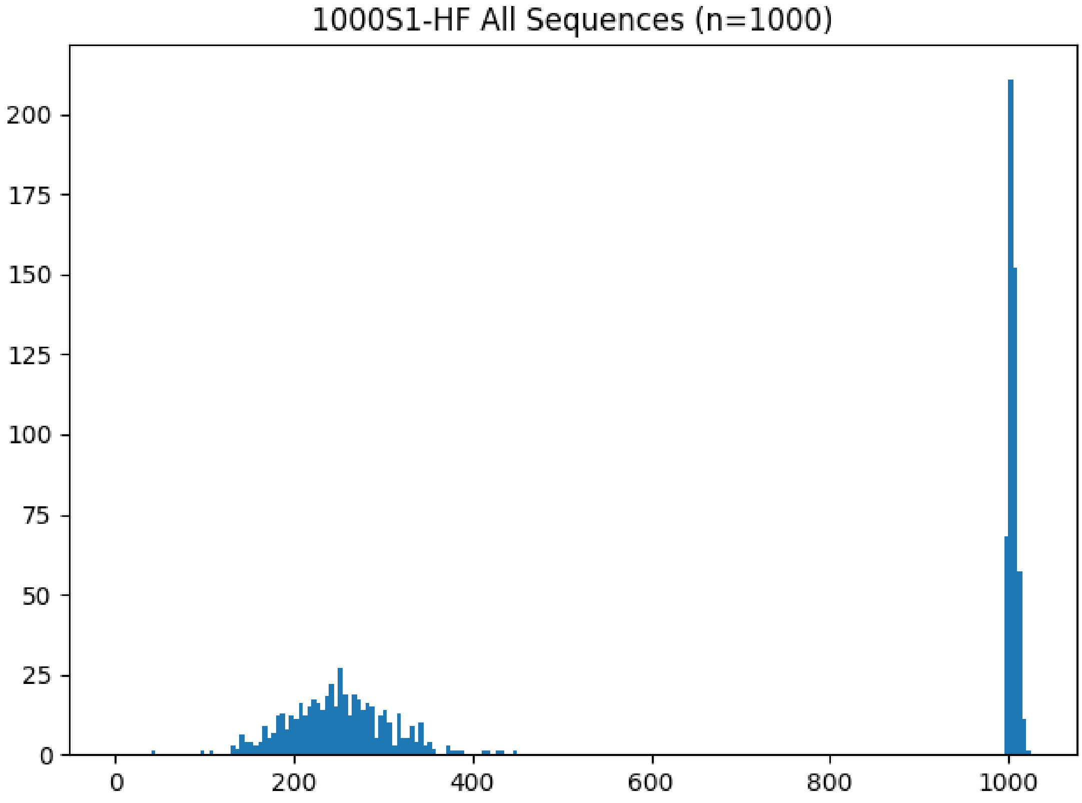

Figure 25: **ROSE 1000S1-HF histogram**

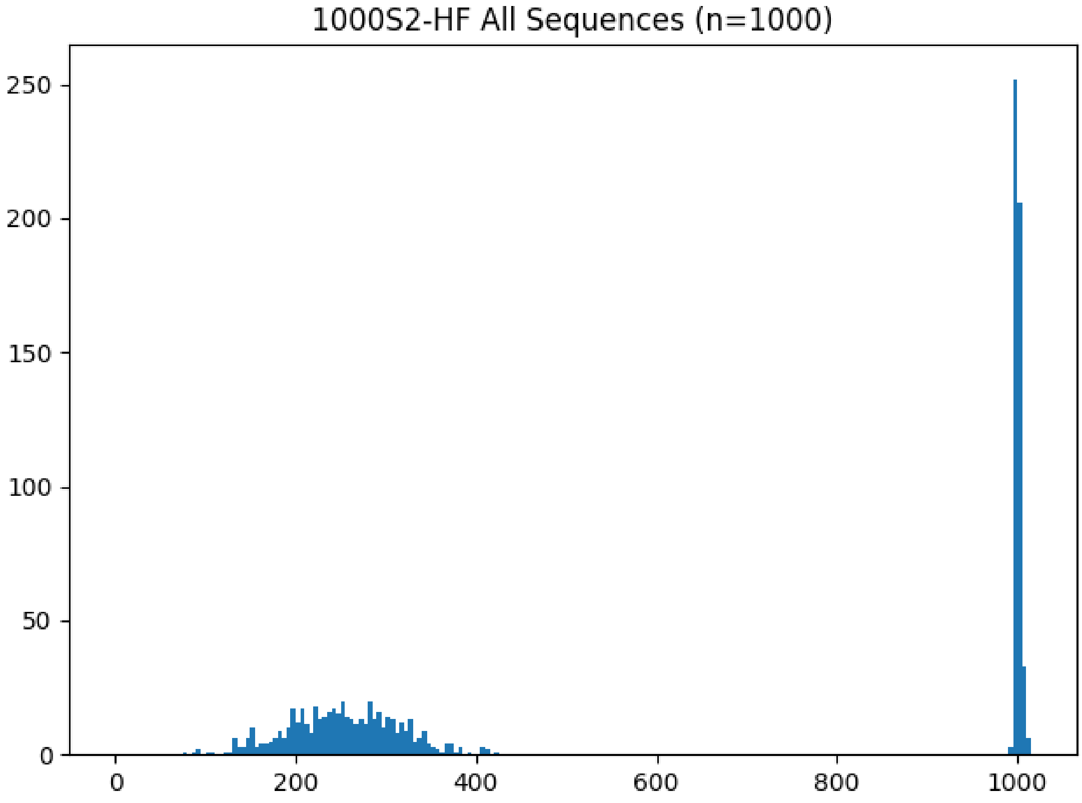

Figure 26: **ROSE 1000S2-HF histogram**

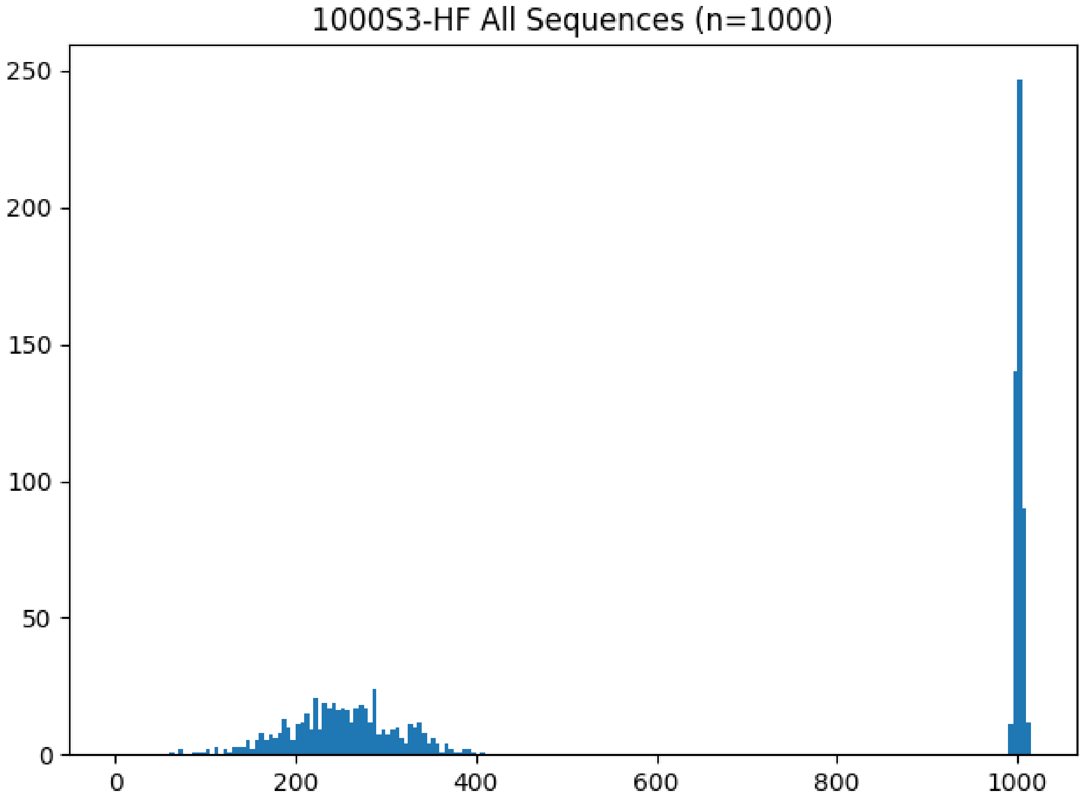

Figure 27: **ROSE 1000S3-HF histogram**

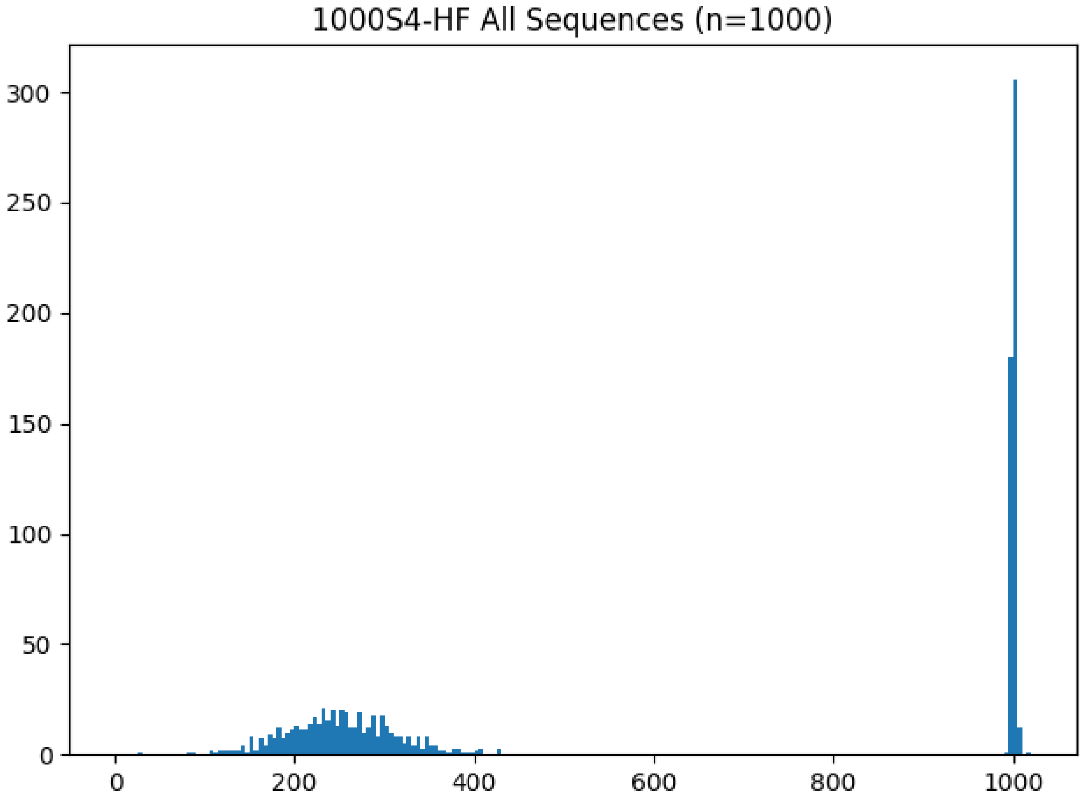

Figure 28: **ROSE 1000S4-HF histogram**

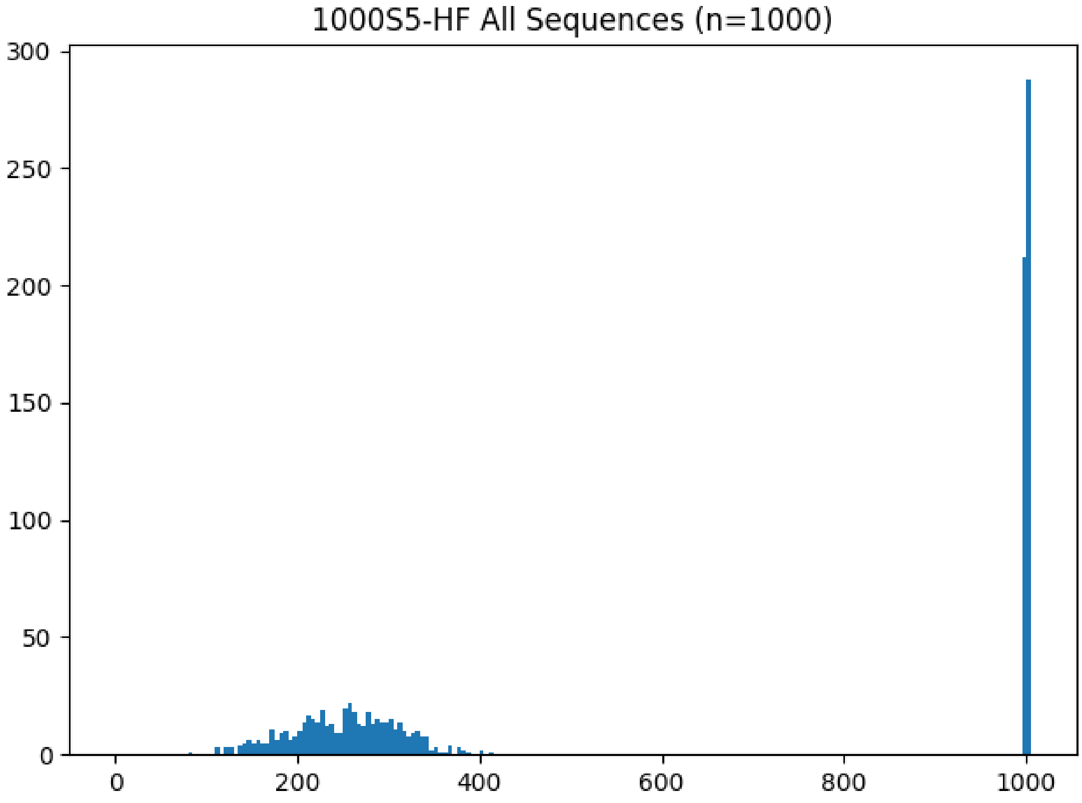

Figure 29: **ROSE 1000S5-HF histogram**

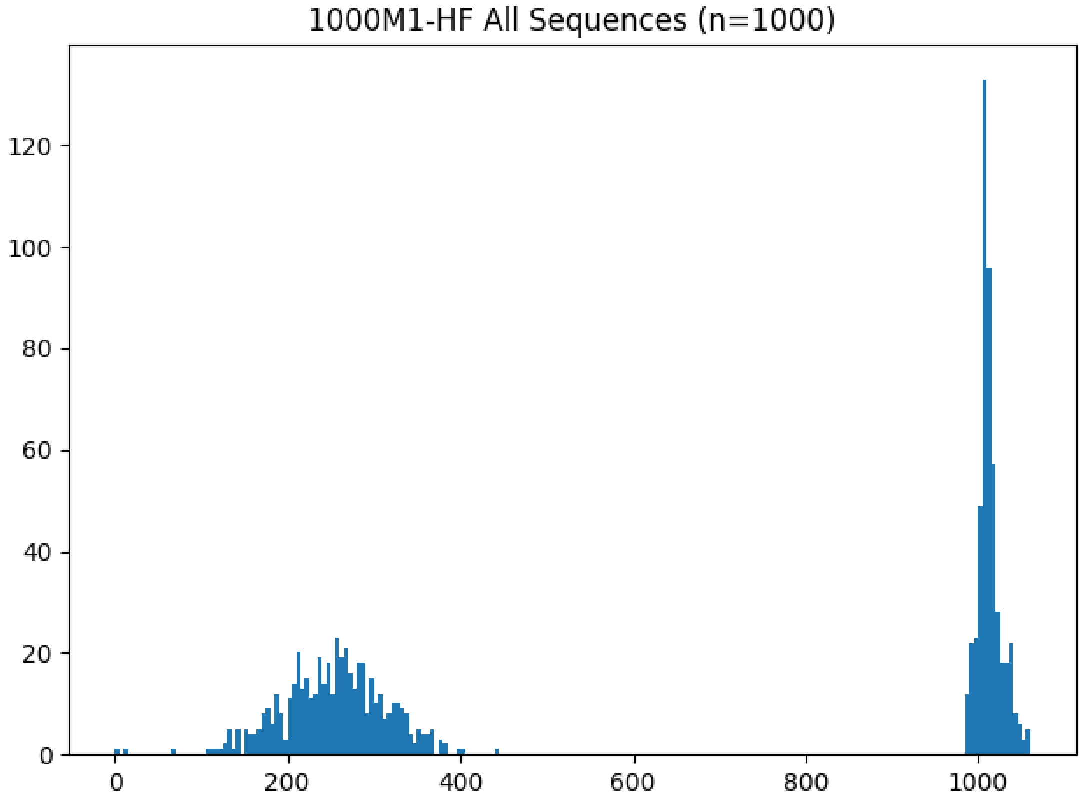

Figure 30: **ROSE 1000M1-HF histogram**

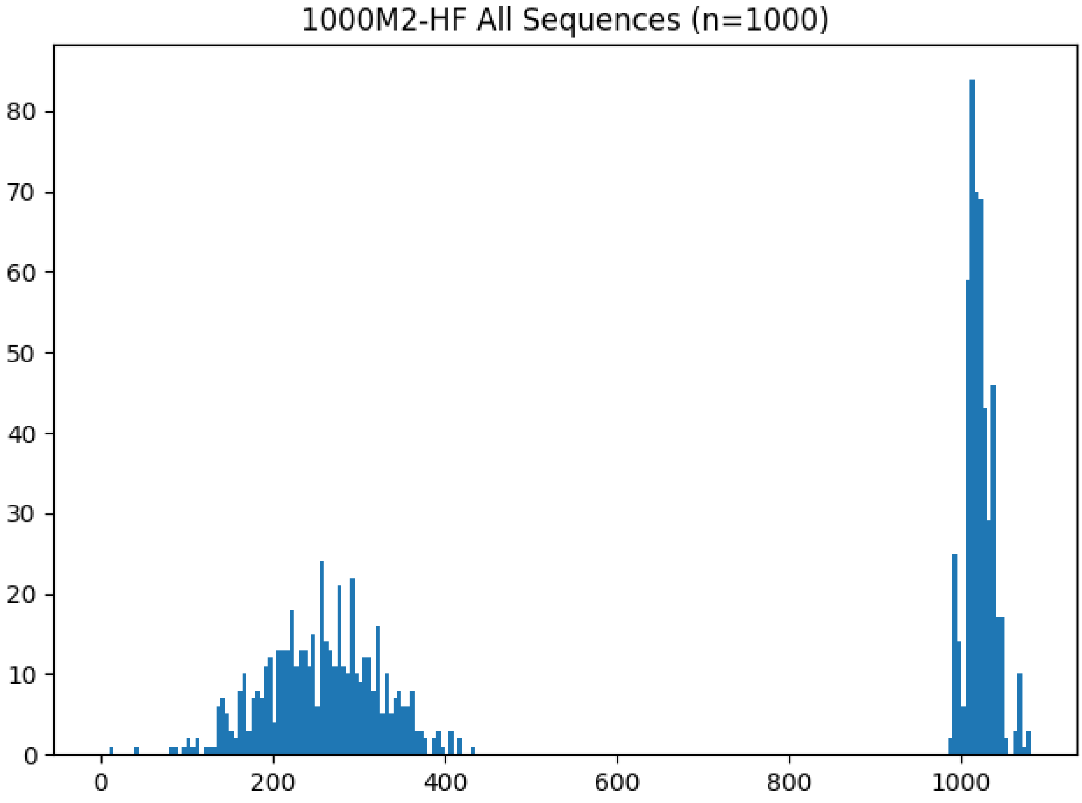

Figure 31: **ROSE 1000M2-HF histogram**

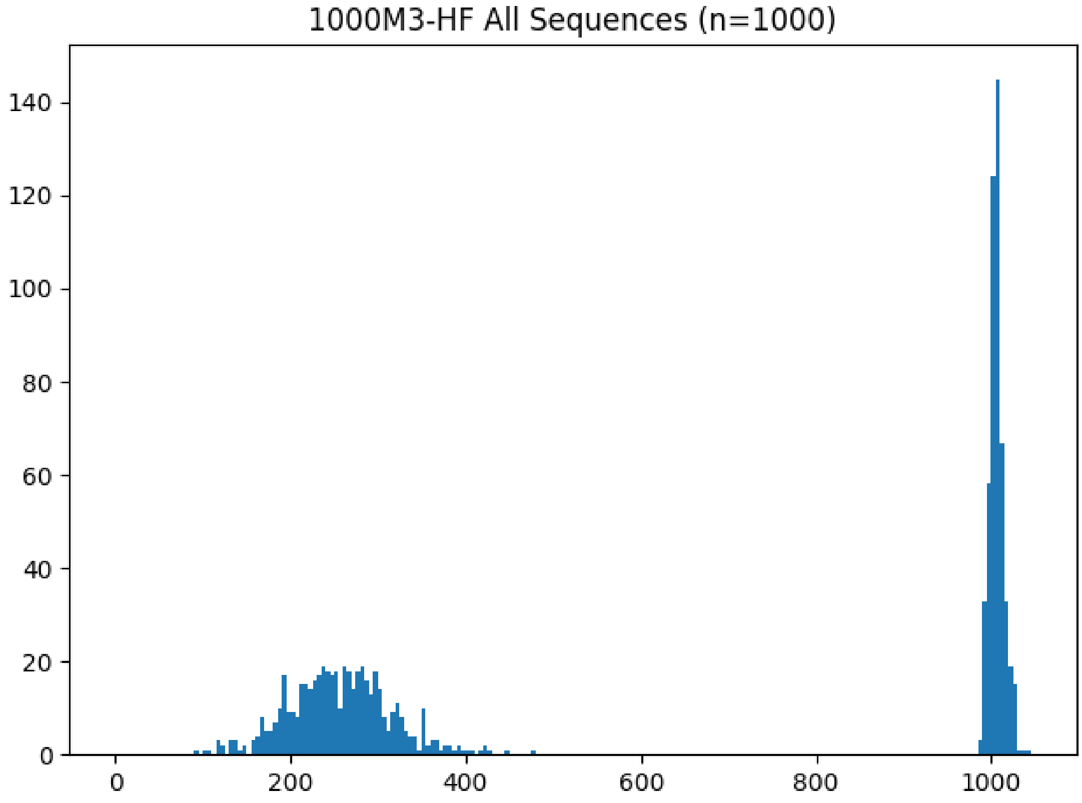

Figure 32: **ROSE 1000M3-HF histogram**

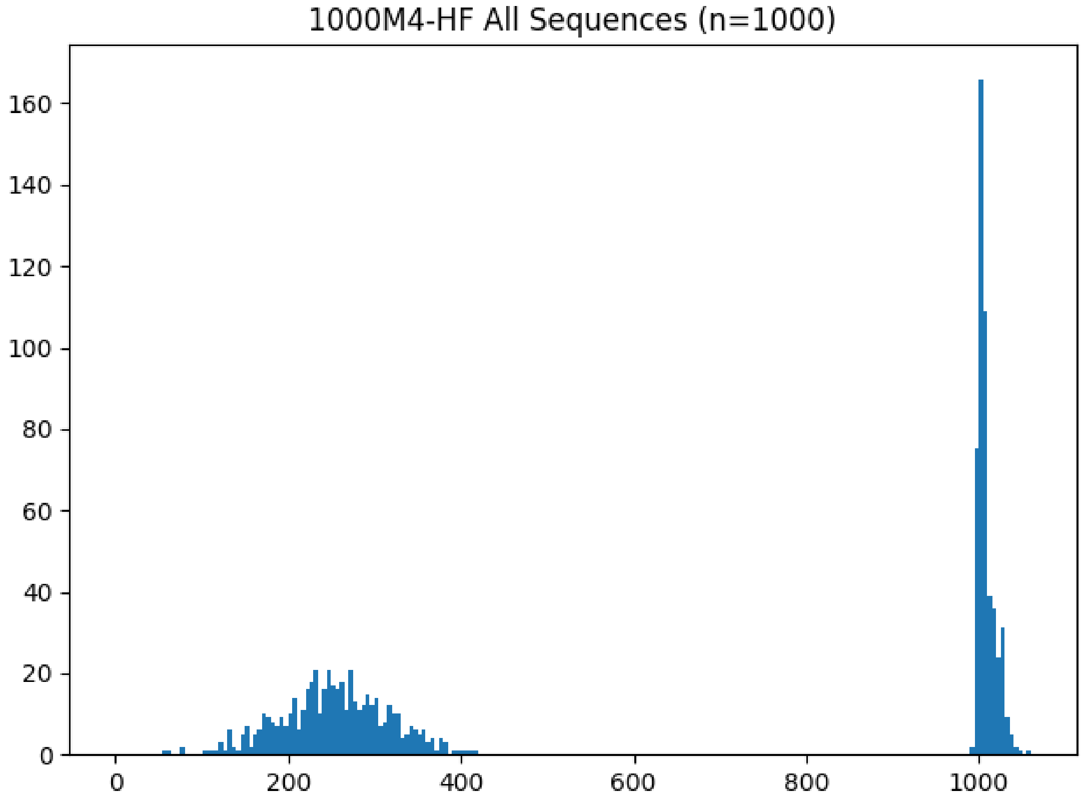

Figure 33: **ROSE 1000M4-HF histogram**

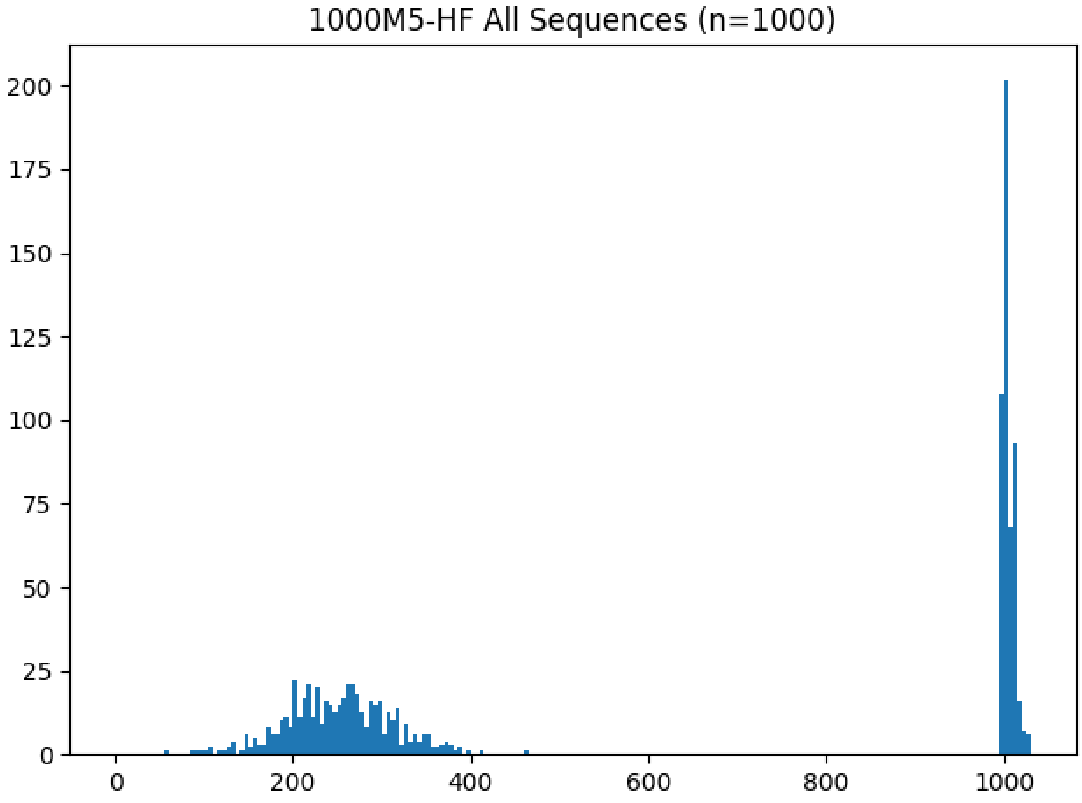

Figure 34: **ROSE 1000M5-HF histogram**

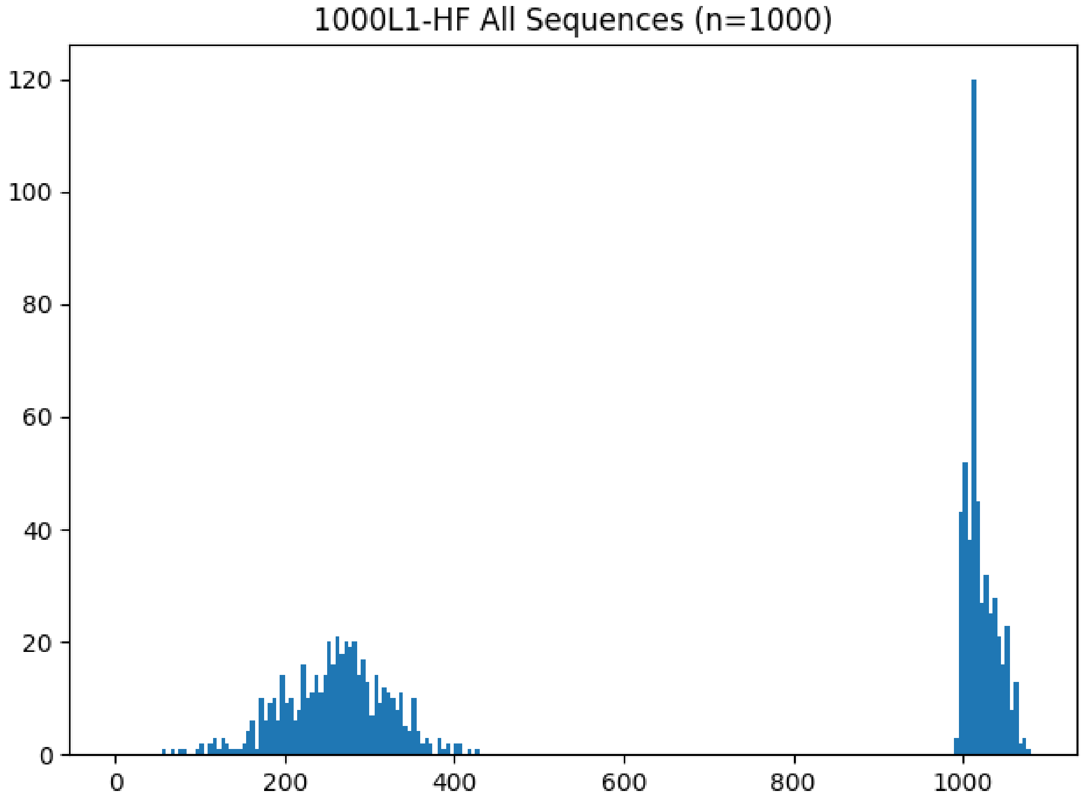

Figure 35: **ROSE 1000L1-HF histogram**

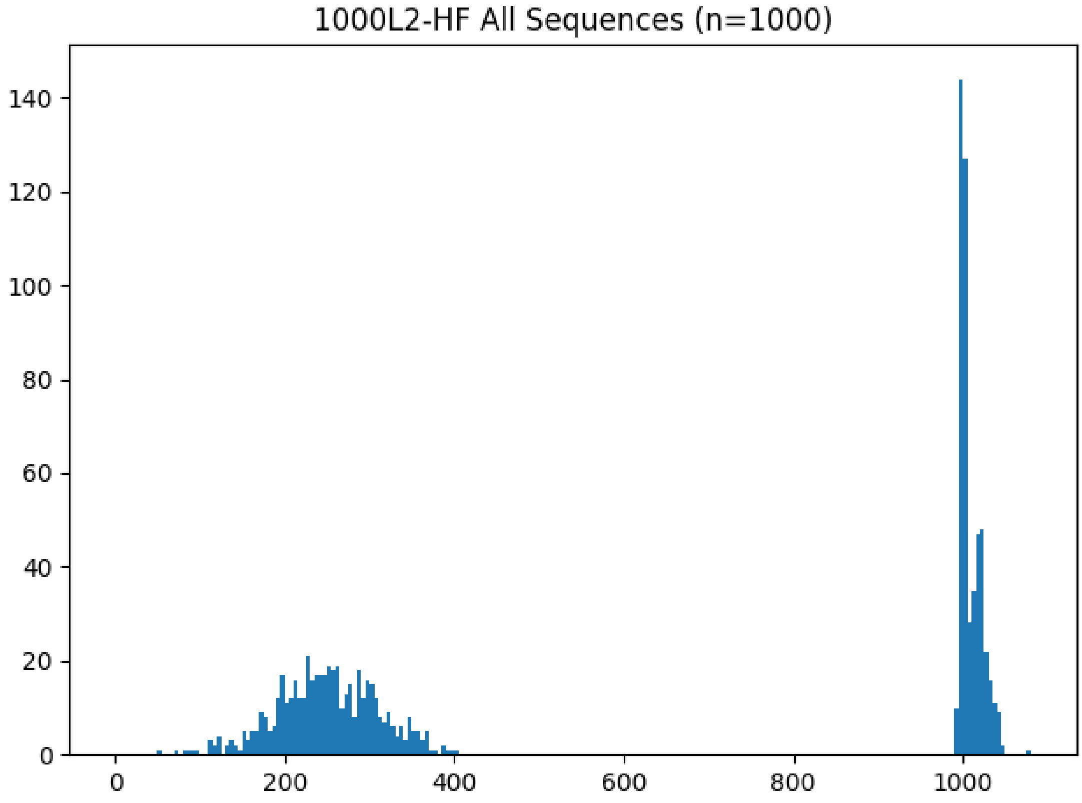

Figure 36: **ROSE 1000L2-HF histogram**

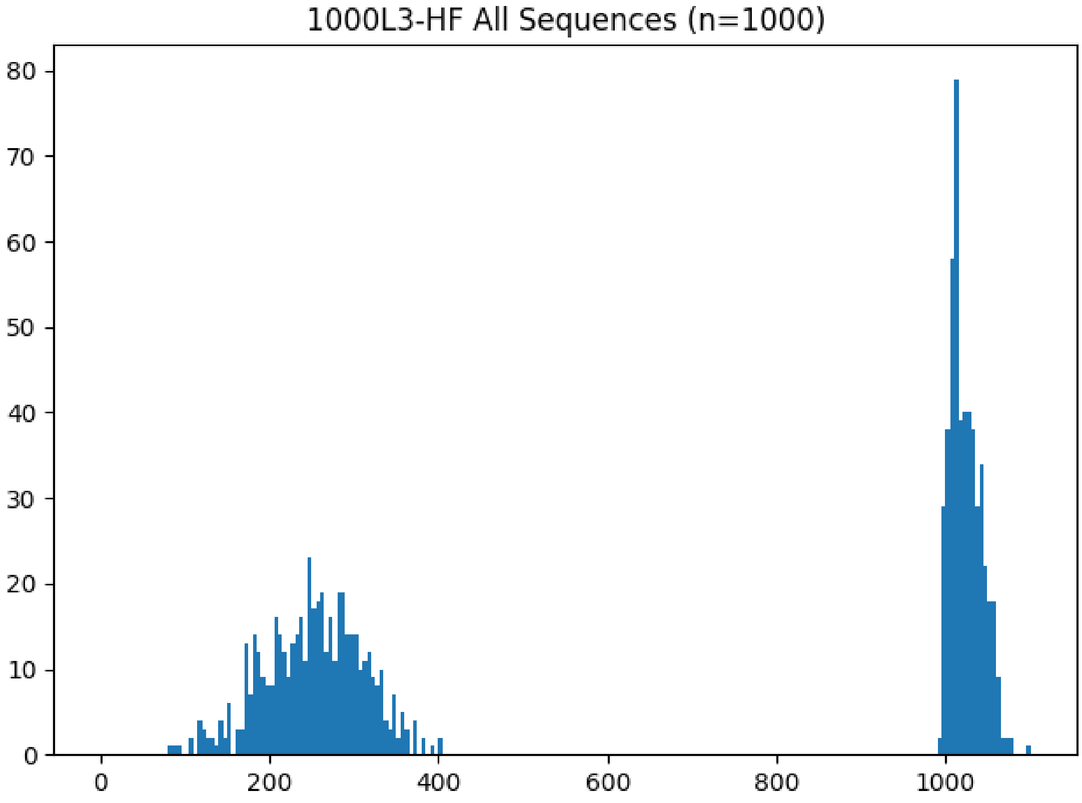

Figure 37: **ROSE 1000L3-HF histogram**

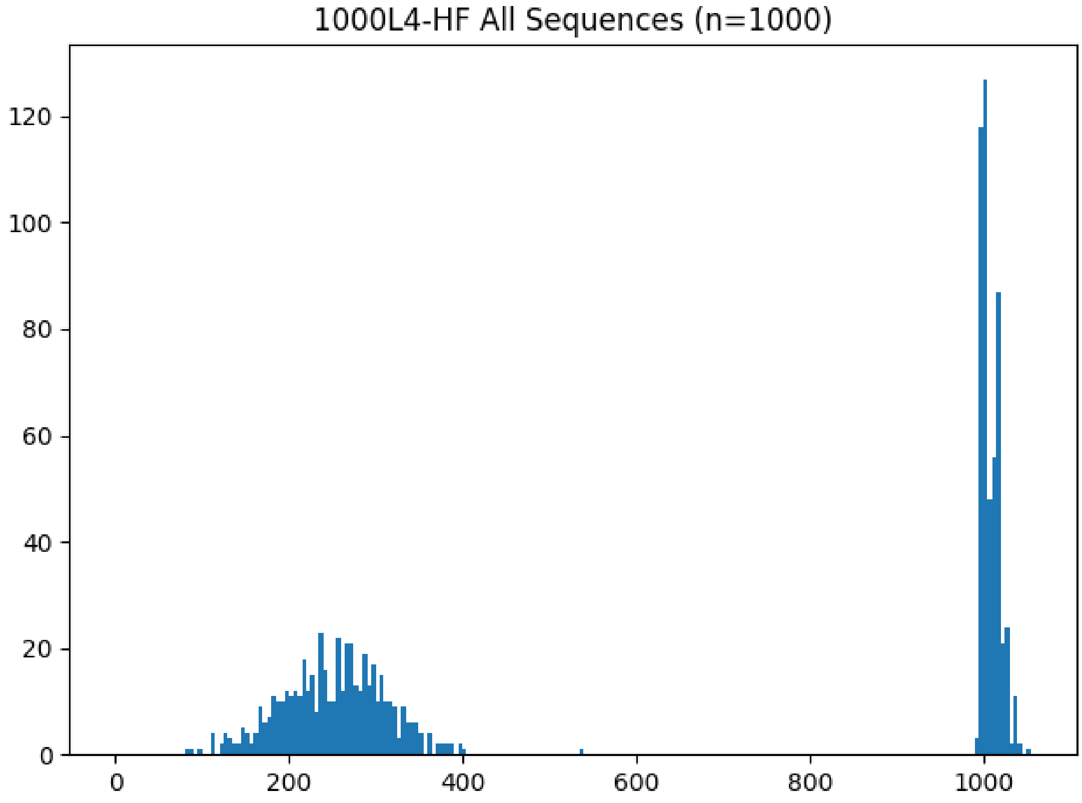

Figure 38: **ROSE 1000L4-HF histogram**

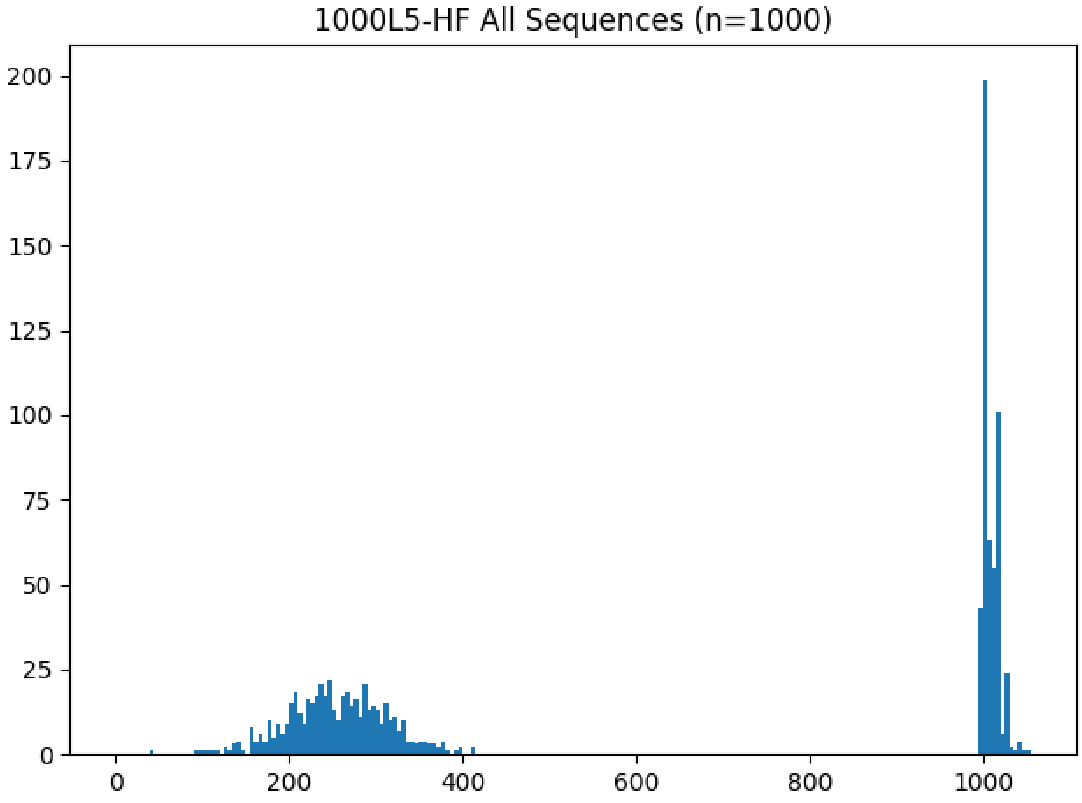

Figure 39: **ROSE 1000L5-HF histogram**

##### 4.5 RNASim Simulated Datasets Sequence Length Histograms

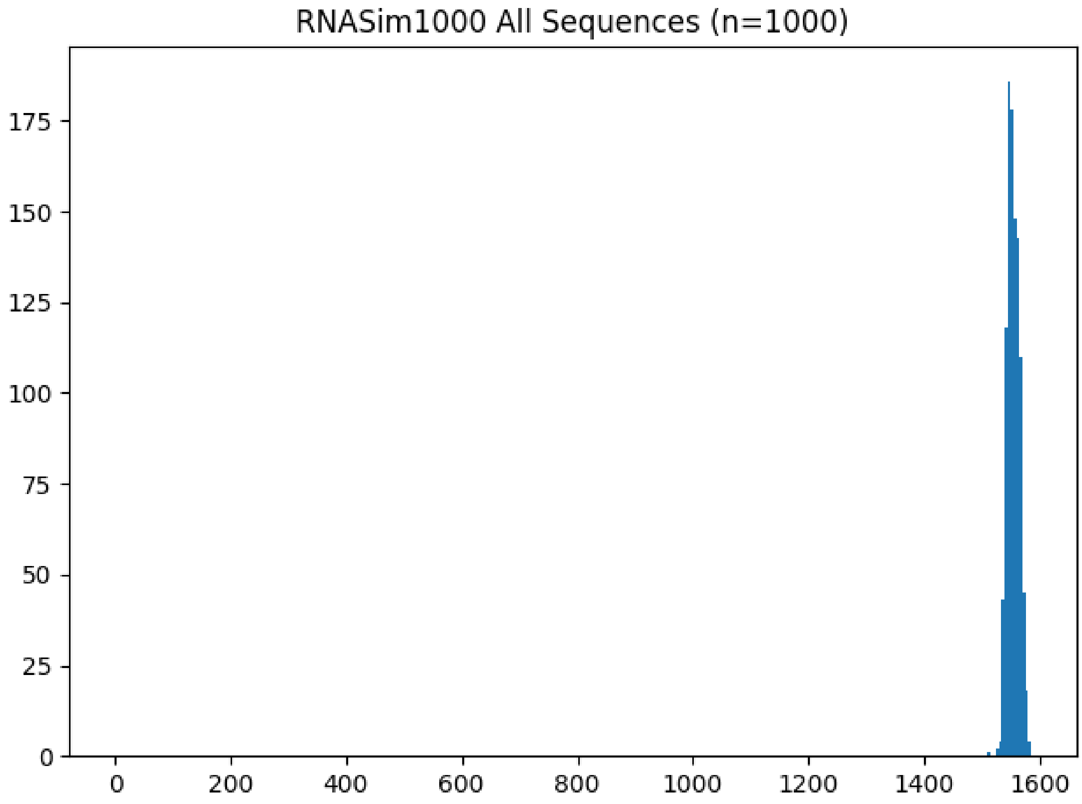

Figure 40: **RNASim1000 histogram**

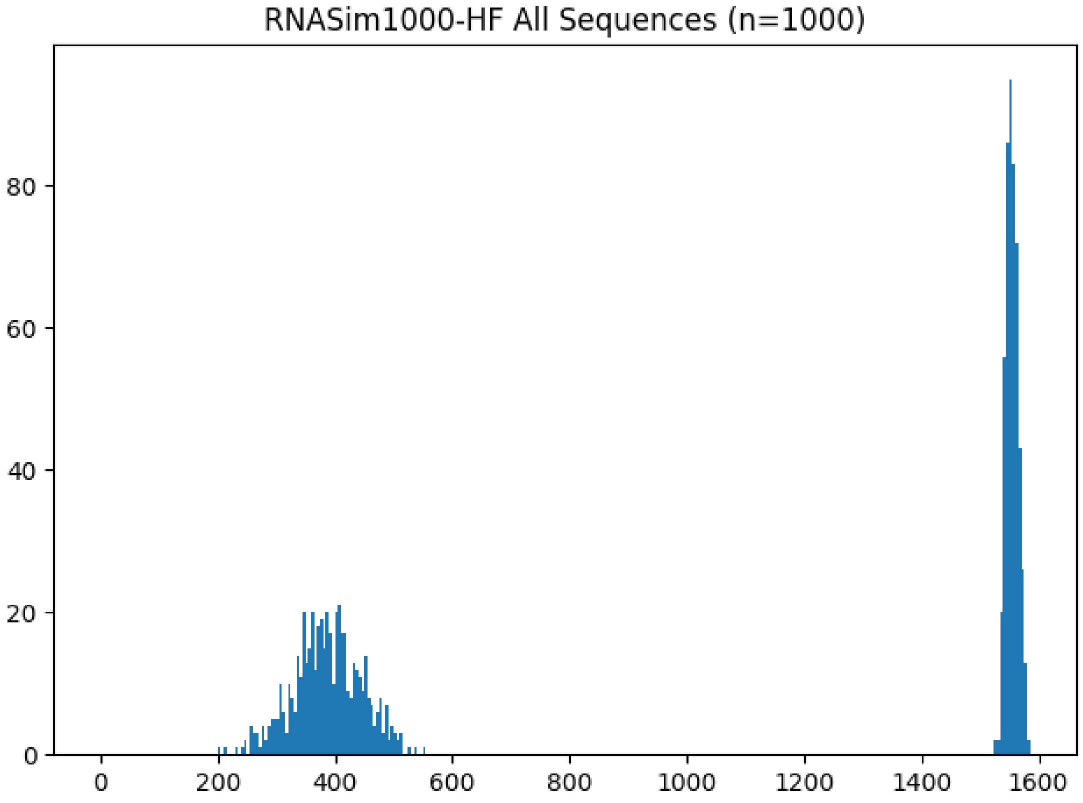

Figure 41: **RNASim1000-HF histogram**

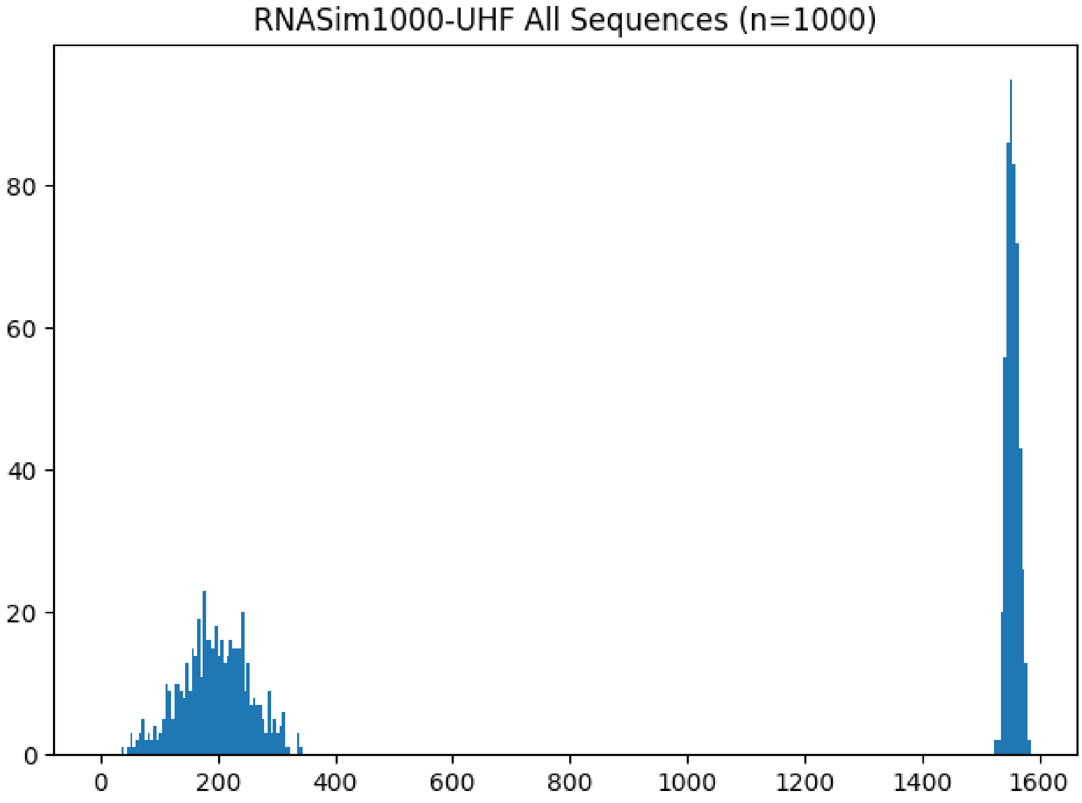

Figure 42: **RNASim1000-UHF histogram**

##### 4.6 CRW Biological Datasets Sequence Length Histograms

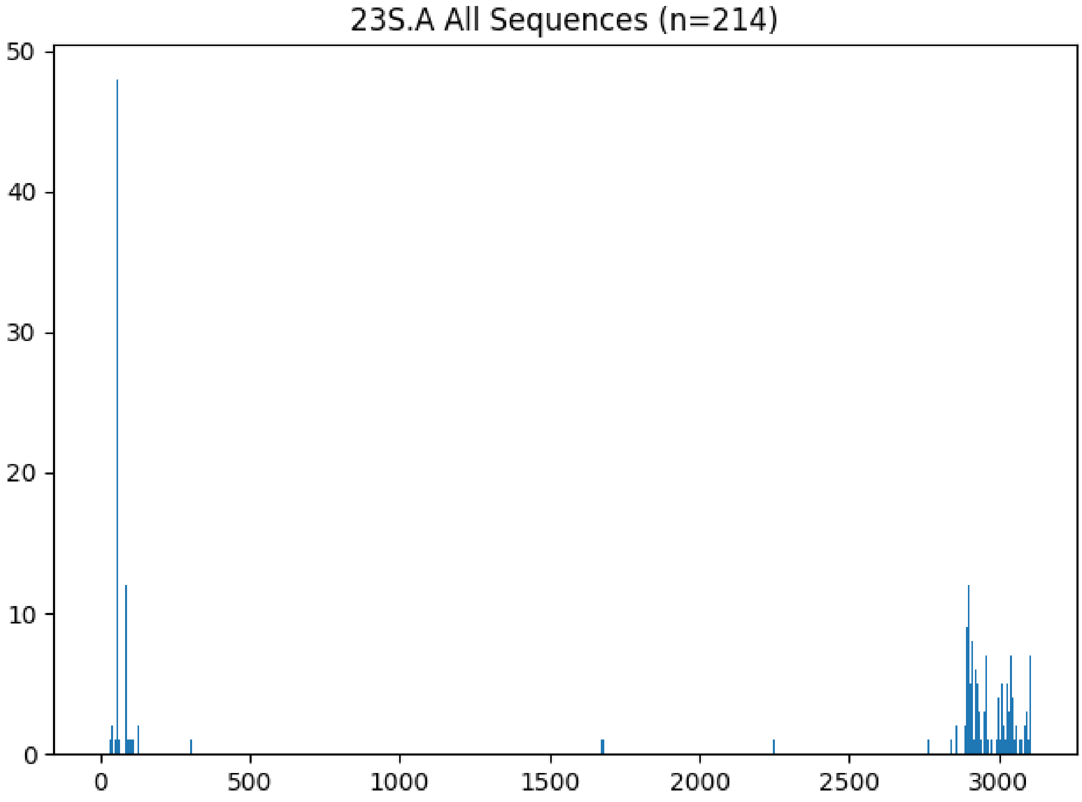

Figure 43: **23S.A histogram**

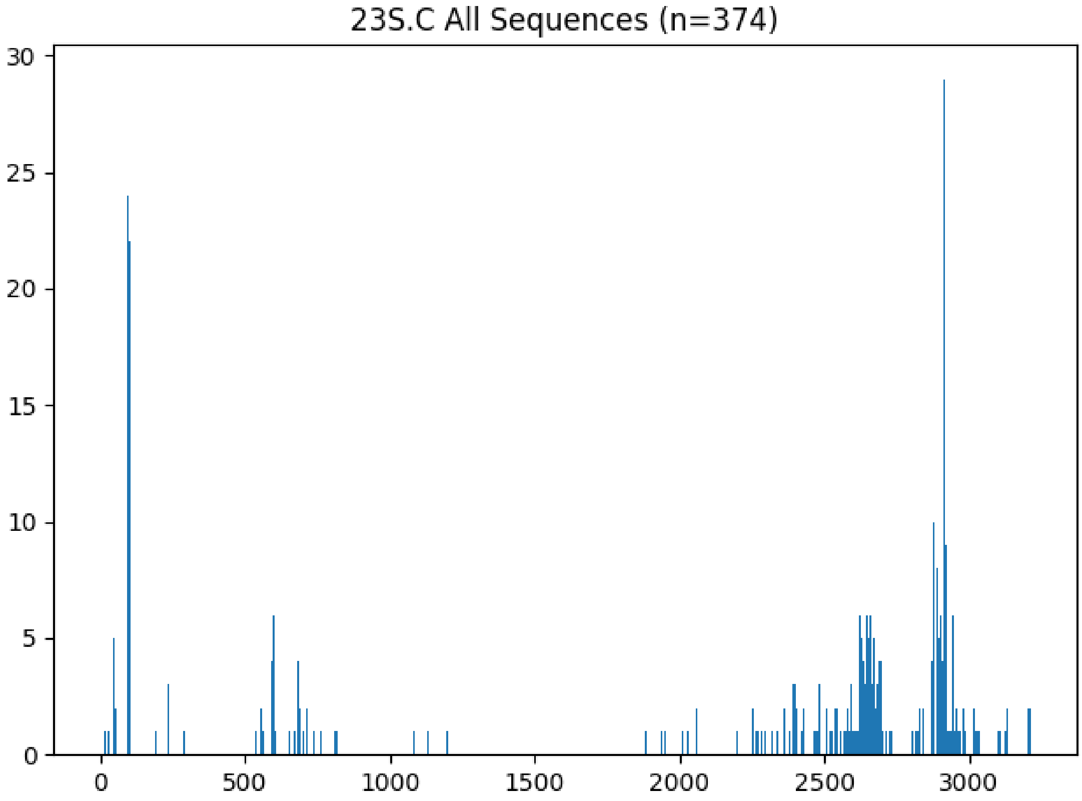

Figure 44: **23S.C histogram**

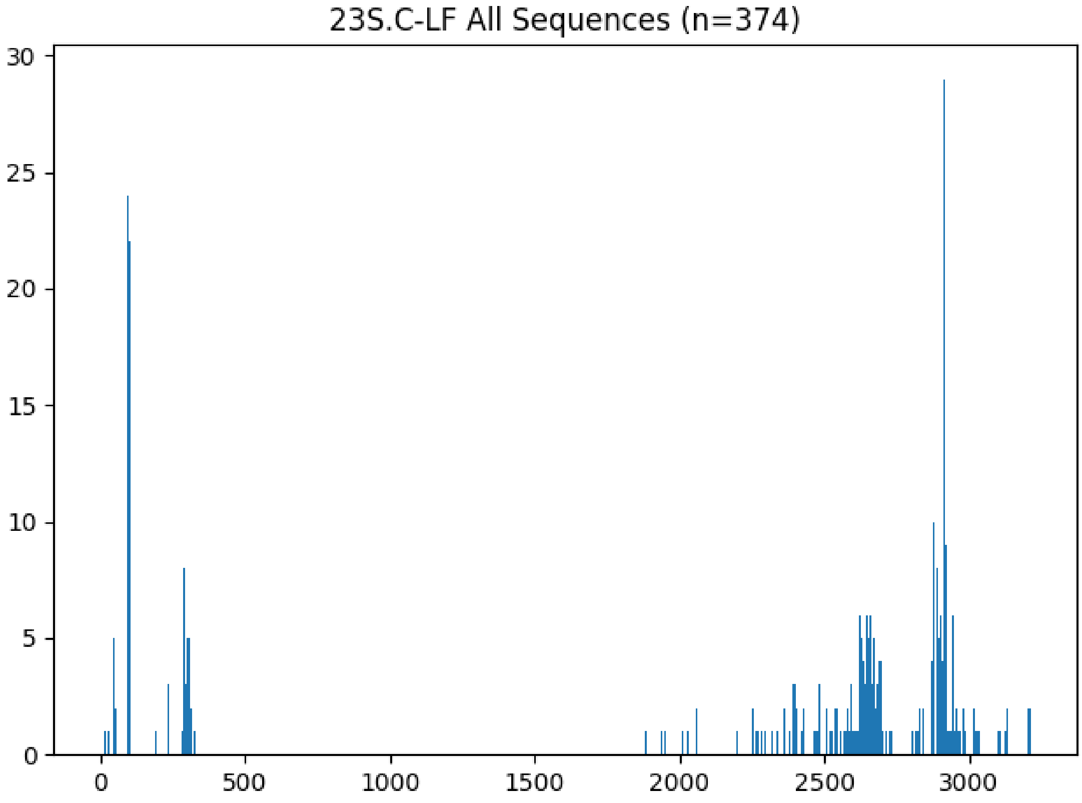

Figure 45: **23S.C-LF histogram**

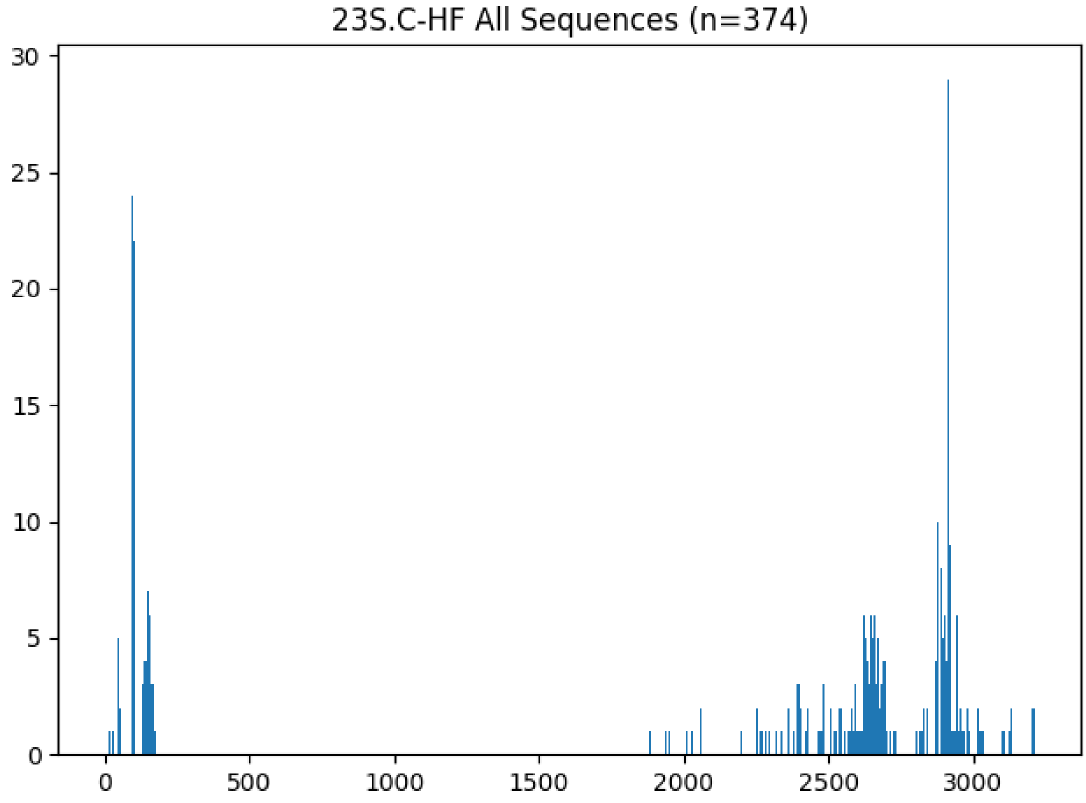

Figure 46: **23S.C-HF histogram**

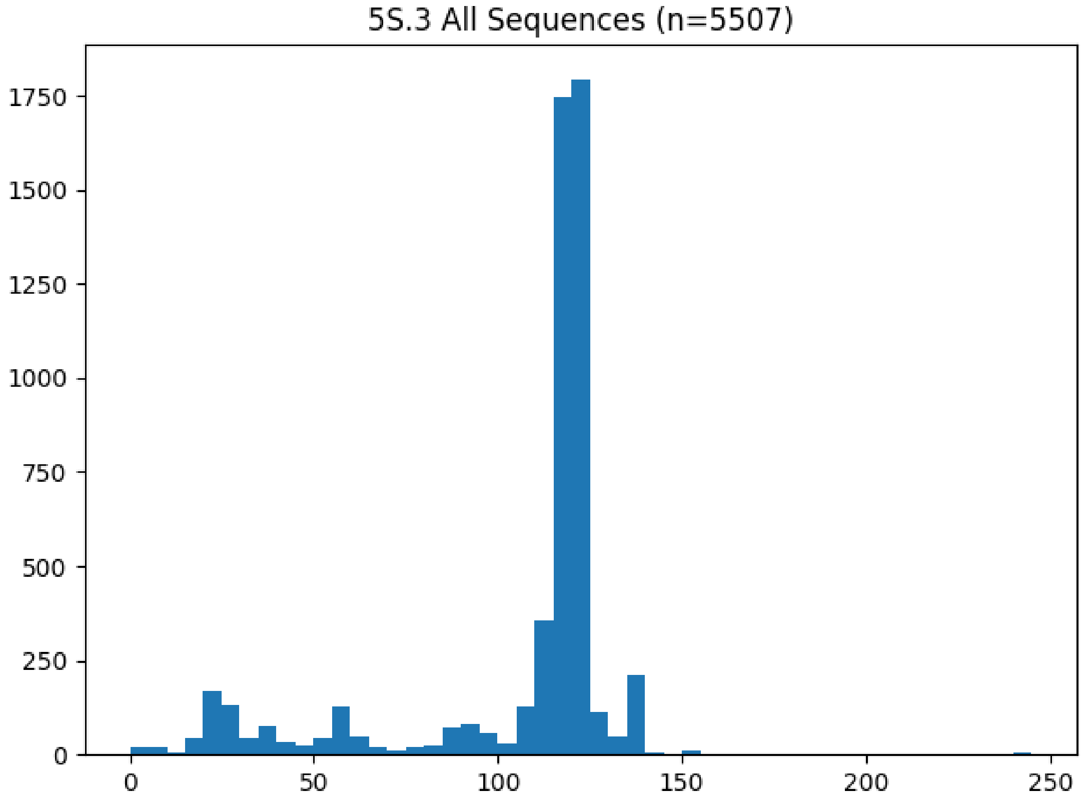

Figure 47: **5S.3 histogram**

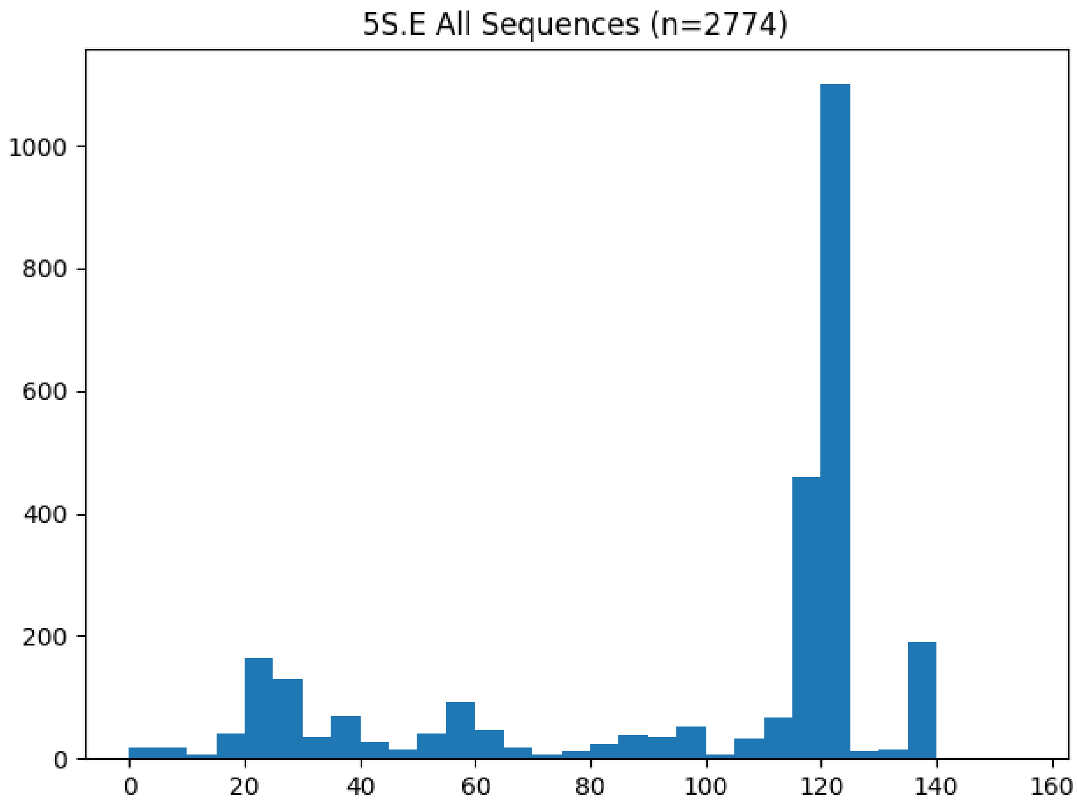

Figure 48: **5S.E histogram**

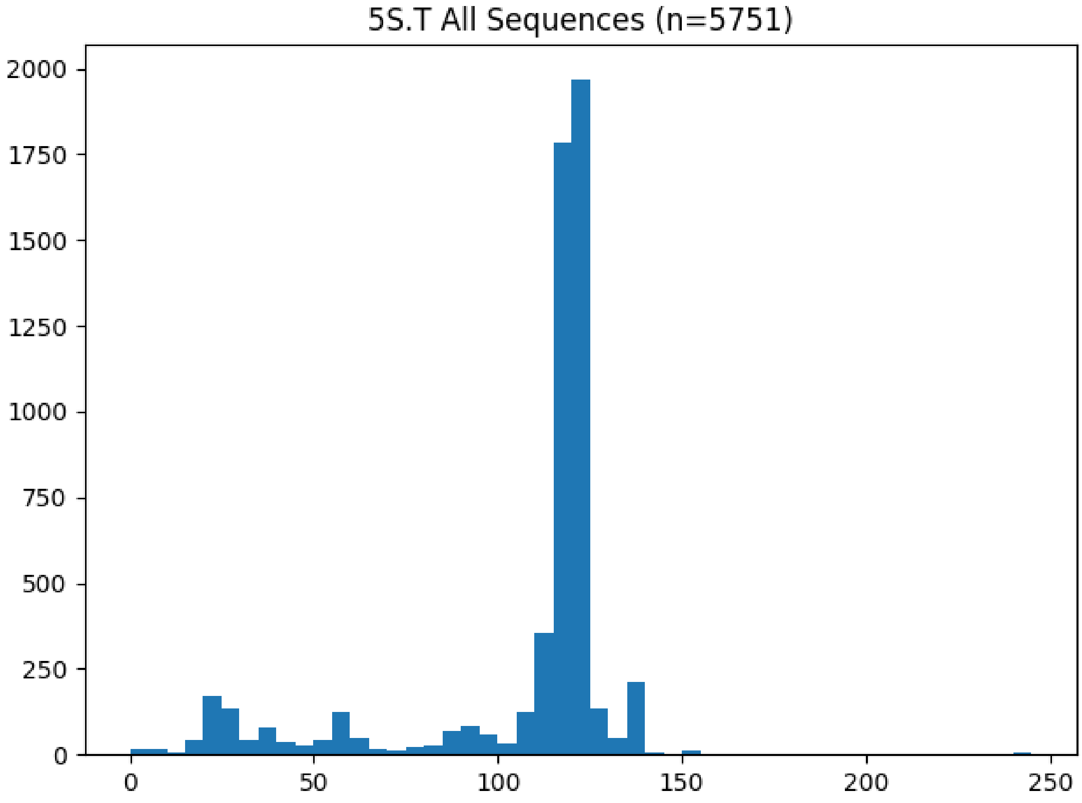

Figure 49: **5S.T histogram**
